# A reduced multicompartment network model of CA1 theta–gamma oscillations under extracellular stimulation

**DOI:** 10.64898/2026.06.22.733913

**Authors:** Maeva Andriantsoamberomanga, Nicolas P. Rougier, Fabien Wagner, Amélie Aussel

## Abstract

Deep brain stimulation has demonstrated its therapeutic potential in modulating pathological oscillations associated with Parkinson’s disease and epilepsy. However, its efficacy in treating disrupted theta-gamma phase-amplitude coupling seen in memory-related disorders, such as Alzheimer’s disease, remains poorly understood. While recent studies have targeted the entorhinal-hippocampal circuit, results remain inconsistent. This discrepancy stems from a lack of mechanistic understanding regarding how stimulation protocols affect this circuit. In this work, we present a reduced multicompartment model of the hippocampal CA1 area that reproduces theta-nested gamma oscillations characteristic of healthy neural activity during memory performance. The model comprises pyramidal, basket and OLM cells with simplified morphologies. We also incorporated CA3-to-CA1 axonal projections, providing a foundational framework for studying how stimulation-induced recruitment of afferent pathways modulates CA1 dynamics. By balancing computational efficiency with anatomical accuracy, our model enables systematic investigation of the effects of electrode placement and orientation, as well as stimulation amplitude and frequency on CA1 neural activity. We demonstrate that the excitatory response in CA1 is primarily driven by the recruitment of Schaffer collateral projections. Overall, this work provides a computationally efficient template for exploring diverse stimulation configurations and could be expanded for developing neuromodulatory strategies to restore physiological network dynamics.

**Author summary:** Deep brain stimulation has shown success in treating Parkinson’s disease by suppressing abnormal neural activity responsible for movement disorders. However, when applied to memory-related pathologies, such as Alzheimer’s disease, the therapeutic outcomes remain unpredictable, ranging from cognitive improvement to impairment. This discrepancy highlights a critical gap in our understanding of how stimulation protocols interact with neural dynamics of the targeted circuits. To address this, we developed a computationally efficient model of the hippocampus, which is involved in memory processes, in order to understand how deep brain stimulation might influence its activity. Our model maintains enough biological accuracy to capture essential memory-related neural activity while remaining lightweight enough for rapid execution and systematic exploration of different protocols. This computational efficiency allowed us to conduct systematic investigations of several stimulation configurations to study their effects on hippocampal dynamics. Overall, this model could provide a useful and computationally cost-efficient tool for exploring the mechanisms of deep brain stimulation and help optimize stimulation protocols aimed at alleviating memory disorders.

## Introduction

The hippocampus plays a central role in episodic memory by processing and integrating information originating from distributed neocortical regions [1, 2]. Information processing and communication is supported by neural oscillations, which coordinate the activity of neurons across hippocampal areas. In particular, theta (4-12 Hz) and gamma (30-100 Hz) rhythms are known to play a role in memory-related operations, including encoding and retrieval of information [3, 4]. Notably, phase-amplitude coupling (PAC) of theta and gamma oscillations, during which the phase of theta oscillations modulates the amplitude of gamma oscillations, has been observed in memory tasks [5, 6]. The strength of theta-gamma PAC is positively correlated with the efficacy of encoding and retrieval, as well as overall mnemonic performance [7–9]. However, these oscillatory patterns are vulnerable to neurodegeneration. An alteration of theta-gamma coupling is observed in the hippocampus during the early stage of Alzheimer’s disease [10–13], making theta-gamma coupling a potential early biomarker of the disease.

Given this fundamental role of rhythmic synchrony, interventions that aim at restoring these oscillatory patterns have gained significant interest for their therapeutic potential. In particular, the effects of deep brain stimulation (DBS) of the entorhinal-hippocampal circuit have been tested in epileptic patients across various studies [14–21], leading to inconsistent findings as highlighted by recent reviews [22–25]. Stimulation of entorhinal cortex or hippocampus has yielded results ranging from enhancement [14–16] and impairment [17, 18] to no significant effect on memory performance [19]. This discrepancy in results may originate from variations in stimulation protocols. For instance, Titiz and colleagues [15] emphasized the importance of precise anatomical targeting, while Merkow and colleagues [18] demonstrated that effects are time-dependent. Furthermore, recent works have suggested that closed-loop stimulation could improve memory performance, whereas random stimulation may impair it [20, 21]. These variations in parameters, including spatial, temporal, and frequency-based characteristics, present a vast array of stimulation configurations that must be systematically investigated to optimize therapeutic efficacy.

Computational modeling provides an essential methodology for elucidating the underlying mechanisms of neural oscillations and evaluating the impact of neuromodulation on these dynamics. Models span a wide spectrum of biological realism, ranging from abstract mathematical mean-field representations [26, 27] to highly detailed conductance-based models that incorporate complex neuronal morphology [28, 29]. While the latter offer high fidelity, their computational demands can be prohibitive. Other approaches reduce complexity by simplifying or even omitting dendritic morphology [30–32]. Despite these differences in scale, these models converge on the emergence of gamma oscillations from recurrent excitatory-inhibitory interactions or within interconnected inhibitory networks. The origin of theta rhythmicity, however, differs across models. Some models impose theta as an external drive to induce theta-gamma coupling [26, 30, 32], whereas in others, theta emerges endogenously from network dynamics [27, 28]. Although Vardalakis and colleagues [32] have begun investigating the effects of neuromodulation on theta-gamma coupling, they modeled the electrical stimulation as an intracellular current applied equally to neurons in the targeted area of stimulation. This simplification neglects the fact that the effects of extracellular electrical stimulation cannot be simply reduced to an intracellular current as they are governed by the second spatial derivative of the extracellular potential along the cell membrane [33, 34].

Computational models specifically addressing hippocampal DBS remain sparse and often incorporate high-fidelity electrode geometries within large-scale networks, leading to high computational costs [35, 36]. For instance, Bingham and colleagues [35], as well as Farzad and colleagues [36], developed computational models of the dentate gyrus comprised of granule cells with explicit inputs from the perforant path. By reconstructing the anatomy of the dentate gyrus from a mouse atlas, they accurately captured tissue resistivity heterogeneity to identify optimal stimulation parameters for network activation. However, while these high-fidelity models allow for precise prediction, they are computationally demanding and do not account for the effects of stimulation on network oscillations. In this work, we present a computationally efficient model to study the impact of extracellular stimulation on theta-gamma oscillations within a model of the human CA1 network. While simpler models treating the electrode as an ideal point source have been successfully applied to the nerve fibers [37–39], to our knowledge, such an approach has not yet been implemented to study hippocampal theta-gamma dynamics.

To enable systematic investigation of various stimulation protocols, we aim at designing a reduced model of CA1 that offers the same functional and morphological properties as a large-scale model. However, this reduction is far from being a linear process regarding population size, parameters and morphological properties. More precisely, the reduced model must exhibit known properties (theta-gamma dynamics) while offering the possibility of investigating extracellular stimulation in a reasonable time. Extracellular stimulation produces a gradient of extracellular potential along the neuron. These spatial differences in extracellular voltage induce an axial current flow within the intracellular space. Such effects cannot be captured by single-compartment neuron models. Furthermore, different parts of a neuron respond differently to electrical stimulation, highlighting the need for multicompartment neuron models to accurately represent extracellular stimulation [33, 34]. While the hippocampus exhibits a vast diversity of interneurons [40, 41], previous studies have suggested the contribution of both parvalbumin-expressing (PV+) basket cells and oriens lacunosum moleculare (OLM) cells in theta-gamma PAC [29, 42, 43]. Therefore, our network focuses on reduced models of basket, OLM and pyramidal cells. As stimulation preferably activates axonal terminals projecting to a region rather than the local cells [44, 45], we explicitly represent Schaffer collaterals projecting from the CA3 area of the hippocampus to CA1 as myelinated axons [46, 47]. Our reduced model exhibits theta-nested gamma oscillations while remaining computationally tractable on a portable computer.

Using this model, we systematically investigated how variations in stimulation amplitude, frequency, and electrode configuration affect the network activity. In addition, we explored the contribution of Schaffer collaterals to the network’s response by comparing stimulation-induced dynamics in both intact and collateral-ablated models. Our results underscore the importance of electrode location and orientation on cell recruitment. Furthermore, we demonstrated that Schaffer collaterals exert a modulatory effect on the network as stimulation induces an increase of activity in their presence, whereas a partial decrease in activity is observed in their absence.

## Results

### Development and validation of the model

To investigate the effects of extracellular electrical stimulation of CA1, we first built a multicompartment network model of a hippocampal slice that can generate theta-nested gamma oscillations (see Material and methods for details). Our network model is made of pyramidal, basket and OLM cells. Axonal trajectories of Schaffer collaterals emerging from CA3 and synapsing onto CA1 pyramidal neurons were also represented. To facilitate the placement of individual cells, their connections based on distance, and the generation of collateral trajectories, we simplified the topology of the hippocampal formation slice into an S-shape, allowing us to develop an intrinsic coordinate system (Fig. 1).

**Fig 1.**
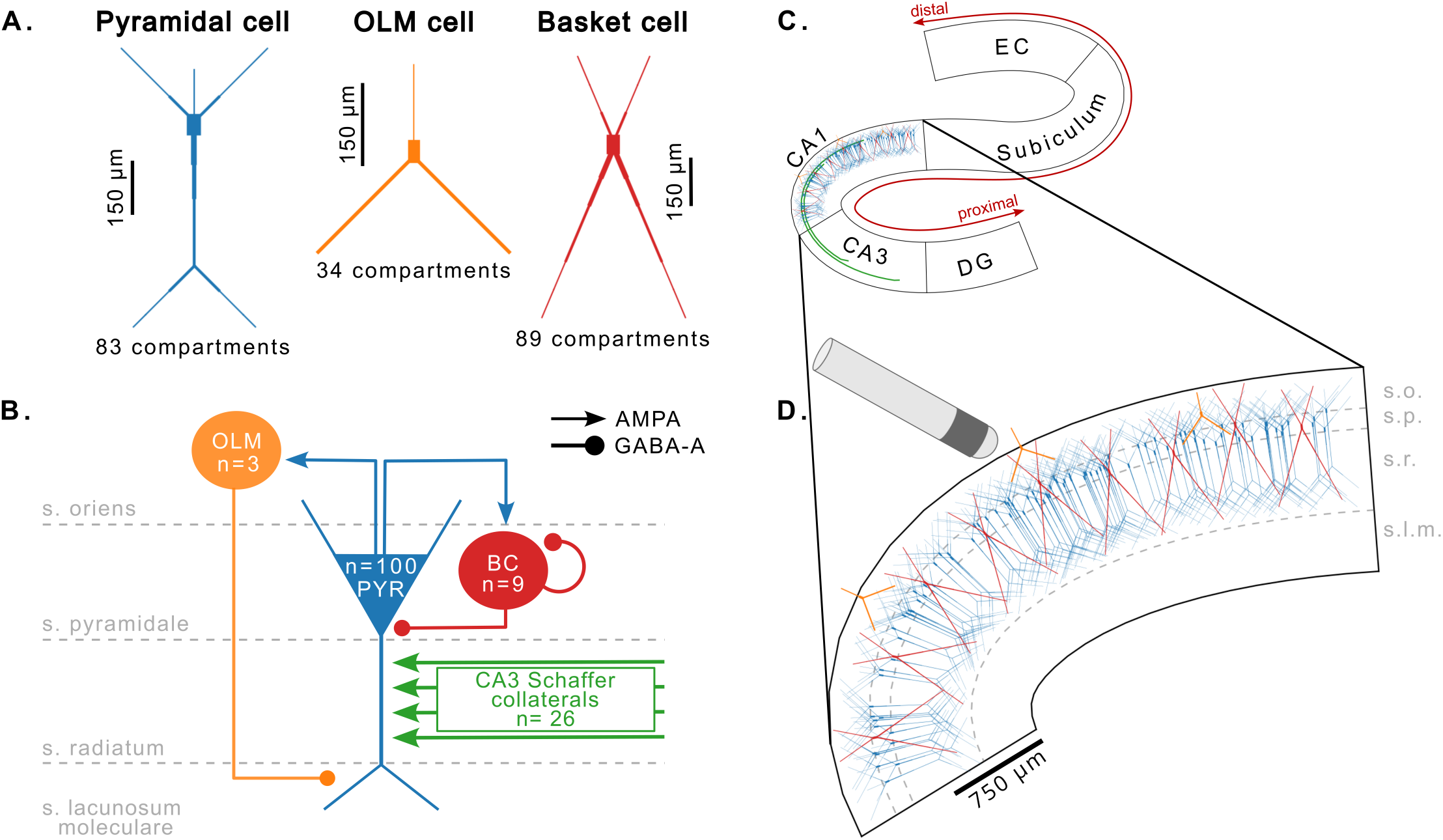
Model overview. **(A)** Multicompartment cells used in the network model. **(B)** Model outline showing the different connections between cell types. The cell types morphologies are simplified for visualization purposes. An external theta input is given to the first nodes of the Schaffer collaterals as an oscillatory intracellular current with an amplitude of 0.093 nA and a frequency of 6 Hz. PYR: pyramidal cells; BC: basket cells. **(C)** General shape of the hippocampal formation slice used for determining the cells placement and the trajectories of myelinated axons projecting from the CA3 area to the CA1 area. EC: entorhinal cortex; DG: dentate gyrus. **(D)** CA1 network showing the placement of cells. The OLM cells soma are placed within the stratum oriens, while the basket cells and the pyramidal cells soma are constrained within the stratum pyramidale. The interneurons are evenly spaced. The principal cells are uniformly placed within the shape of the CA1 area. An extracellular stimulation is given by an electrode located outside of the CA1 area. s.o.: stratum oriens; s.p.: stratum pyramidale; s.r.: stratum radiatum; s.l.m.: stratum lacunosum moleculare

#### Synaptic weight tuning within CA1 enables the generation of gamma rhythms

Assuming that gamma oscillations emerge from the interactions between excitatory and inhibitory neurons, we first tuned the synaptic weights of the CA1 network in the absence of Schaffer collaterals to ensure it would sustain activity in the gamma range (see Fig. 2). An intracellular ramp current with an amplitude increasing from 0 to 1 nanoampere (nA) was injected into the soma of the pyramidal cells. Starting with initial synaptic weights adapted from [32], we performed several 5-second simulations during which the excitatory synaptic weights (*w*_*EOLM*_ and *w*_*EBC*_) were scaled by a factor *k*_*E*_ ranging from 0.1 to 2, and the inhibitory synaptic weights (*w*_*IE*_) by a factor *k*_*I*_ ranging from 0.1 to 1. The selected parameters, *k*_*E*_ = 2 and *k*_*I*_ = 0.1, allowed us to achieve an oscillation frequency of at least 30 Hz in all cell populations under high intracellular stimulation), as depicted by the heatmaps in Fig. 2B, as well as the spectrograms in Fig. 2D. We also chose these values so that we would observe a firing rate that was 2.5 times faster in the basket population than in the pyramidal population (Fig. 2C), which is in accordance with the fact that basket cells have a higher discharge rate than pyramidal cells [48].

**Fig 2.**
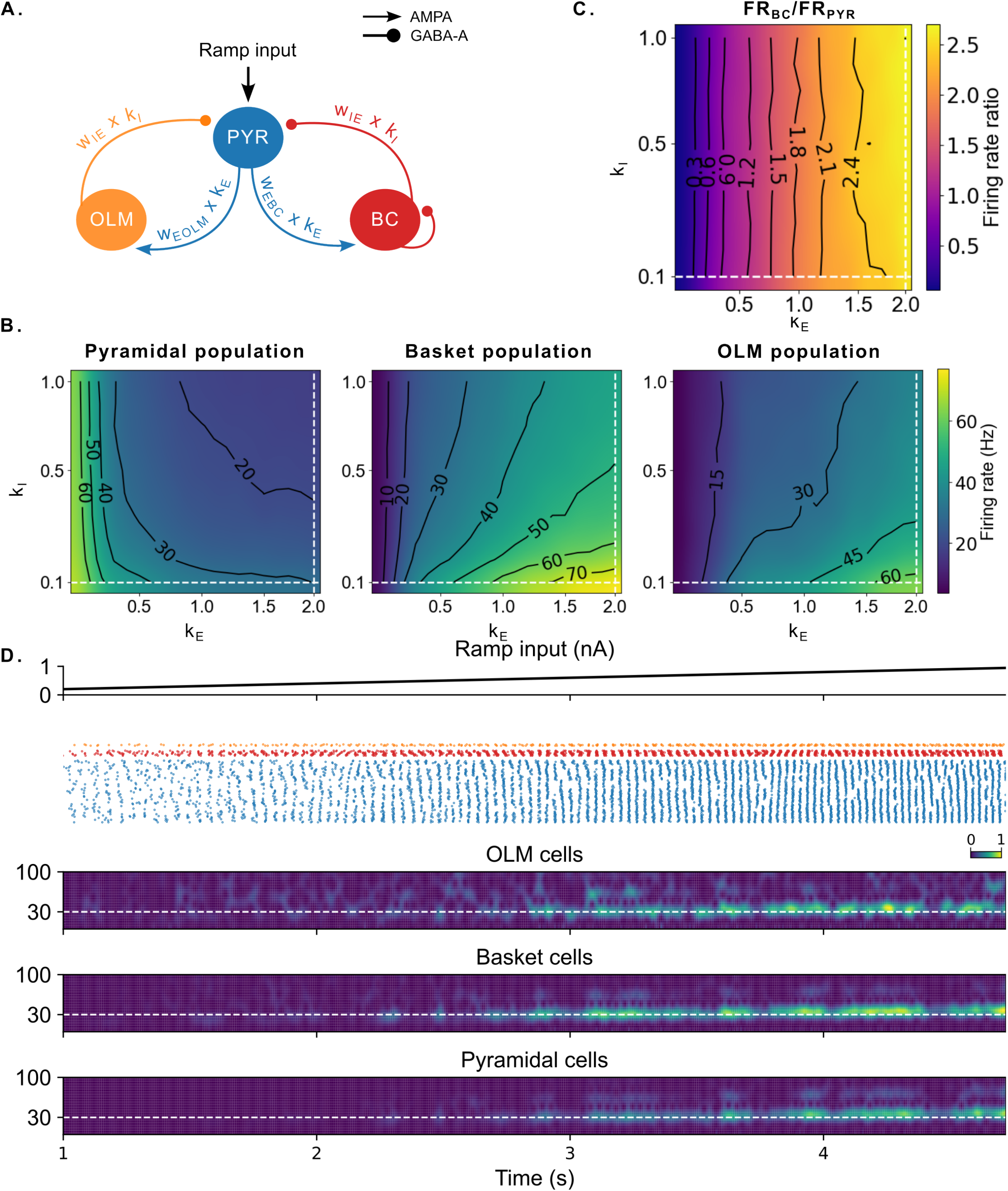
Synaptic weights optimization. **(A)** Schematic representation of the network during the optimization of synaptic weights. OLM-to-pyramidal and basket-to-pyramidal synaptic weights were both initialized with the same value, *w*_*IE*_, adapted from [32]. Excitatory synaptic weights from pyramidal cells to OLM cells (*w*_*EOLM*_) and to basket cells (*w*_*EBC*_) were initialized to elicit slow gamma-frequency activity in both interneuron populations. An intracellular input was injected into the soma of pyramidal neurons in the form of a ramp input, with an amplitude ranging from 0 to 1 nA over a 5-second simulation. Multiple simulations were conducted in which the inhibitory weights (*w*_*IE*_) were multiplied by a factor *k*_*I*_ ranging from 0.1 to 1, while the excitatory weights (*w*_*EOLM*_ and *w*_*EBC*_) were multiplied by a factor *k*_*E*_ ranging from 0.1 to 2.**(B)** Mean oscillatory frequency of each population as a function of the scaling factors, *k*_*E*_ and *k*_*I*_, during the last second of simulation. Dashed lines indicate the mean oscillatory frequency corresponding to the selected parameters (*k*_*I*_ = 0.1 and *k*_*E*_ = 2), which yielded oscillatory frequency of approximately 30 Hz for pyramidal cells, 75 Hz for basket cells, and 60 Hz for OLM cells. **(C)** Ratio of the mean oscillatory frequency between basket cell population and pyramidal cell population during the last second of each simulation. To ensure that basket cells fired at least twice as frequently as pyramidal cells, we computed the ratio of their mean oscillatory frequency over the last second of simulation. The point at the intersection of the dashed lines corresponds to a ratio of approximately 2.5. **(D)** Simulation results obtained with the selected parameters (*k*_*I*_ = 0.1 and *k*_*E*_ = 2), producing gamma-frequency activity across all populations. Under these conditions, interneuron populations exhibited faster activity than pyramidal cells, as indicated by the dashed lines in panels A and B. The top panel shows the ramping current input injected into the soma of pyramidal cells. The second panel shows neuronal spiking activity over time for each population (orange: OLM cells; red: basket cells; blue: pyramidal cells). The last panels display spectrograms of the binned firing rate of each population throughout the simulation. The spectrograms show an increase in power and frequency alongside the ramp input, reaching fundamental oscillatory frequencies between 40 and 50 Hz for all populations.

#### Schaffer collaterals inputs tuning enable theta-gamma oscillations

Once all the CA1 neuronal populations were able to produce oscillations in the gamma frequencies, we added the Schaffer collaterals. 6-Hz oscillatory intracellular currents were injected into the first node of each Schaffer collateral to simulate endogenous theta rhythms emerging from the CA3 area and propagating to the CA1 area. We optimized the amplitude of these oscillatory inputs, as well as the synaptic weights, *w*_*SCA*_, from Schaffer collaterals to pyramidal cells (see Fig. 3). Several 5-second simulations were performed during which the input amplitude, *k*_*amp*_, were varied from 0.02 to 2 nA. The synaptic weights *w*_*SCA*_ were scaled by a factor *k*_*SCA*_ ranging from 2 to 4, starting from a value adapted from the same value as pyramidal-to-basket synaptic weight obtained after optimization. Heatmaps representing the mean oscillatory frequency of pyramidal and basket populations within theta phases were built (Fig. 3B). From these heatmaps, we determined the parameters that induced the strongest gamma activity in both populations. The selected parameters, *k*_*amp*_ = 0.1 nA and *k*_*SCA*_ = 2.9, resulted in a mean oscillatory frequency above 50 Hz in both pyramidal and basket populations nested within the phases of the ongoing theta (Fig. 3B and C).

**Fig 3.**
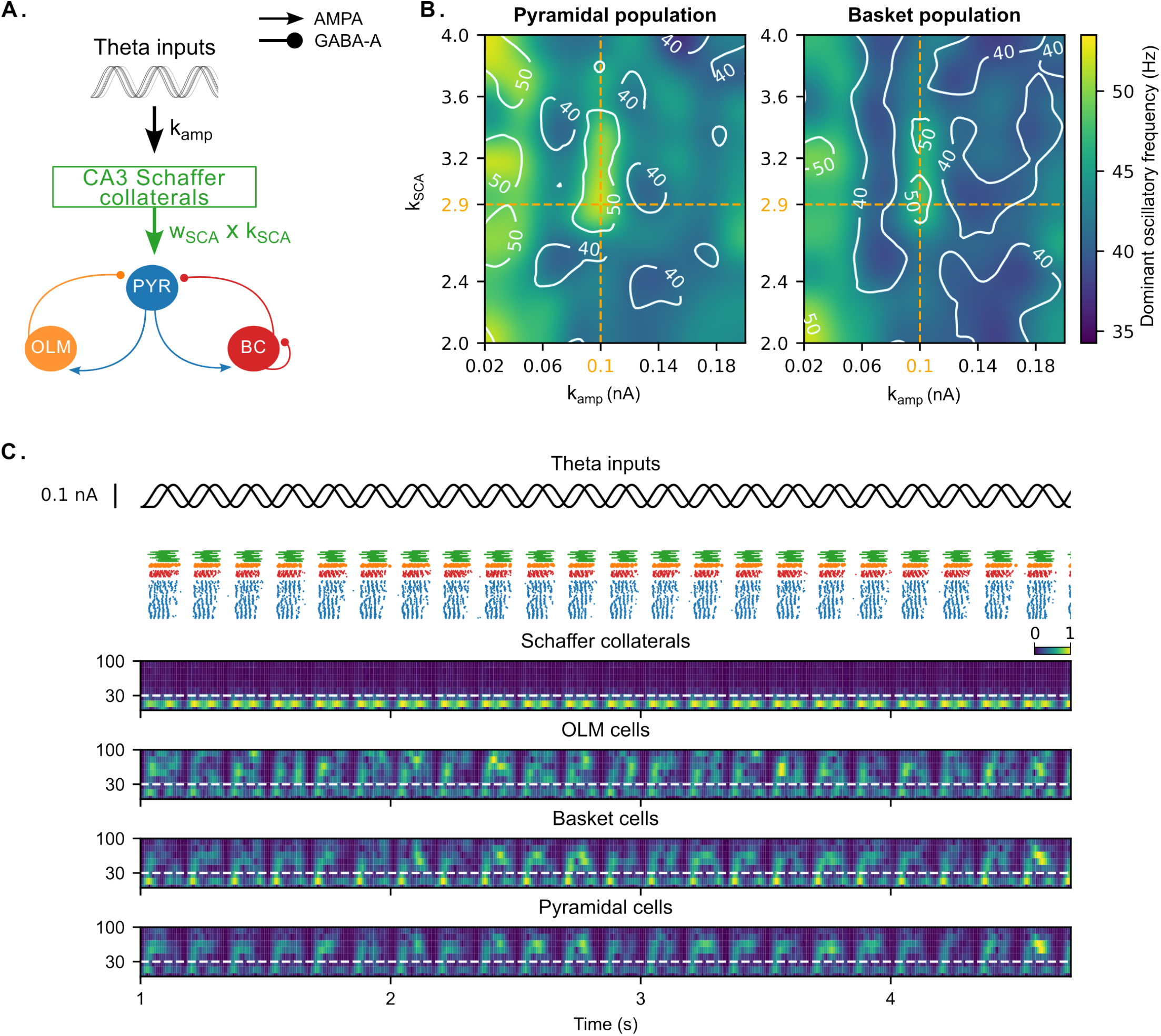
Optimization of input amplitude and Schaffer collaterals synaptic weights. **(A)** Schematic representation of the network during the optimization of input amplitude (*k*_*amp*_) and Schaffer collaterals synaptic weights (*w*_*SCA*_). Theta inputs were injected into the first nodes of the Schaffer collaterals as an oscillatory intracellular current. *w*_*SCA*_ was fixed to the same value as pyramidal-to-basket synaptic weight obtained after optimization. Multiple simulations of 5 seconds were conducted during which *k*_*amp*_ varied from 0.02 to 0.2 nA with a step of 0.02 nA, and *w*_*SCA*_ was multiplied by a factor *k*_*SCA*_ ranging from 2 to 4. **(B)** Mean oscillatory frequencies of pyramidal and basket populations within theta phases as a function of the input amplitude and scaling factor for Schaffer-to-pyramidal weight. The orange dashed lines indicate the selected parameters (*k*_*amp*_ = 0.1 nA and *k*_*SCA*_ = 2.9), which maximizes the oscillatory frequencies of both populations. **(C)** Simulation results obtained with the selected parameters (*k*_*amp*_ = 0.1 nA and *k*_*SCA*_ = 2.9). The first panel shows the theta-oscillatory current injected into the Schaffer collaterals. The second panel shows neuronal spiking activity over time for each population (green: last nodes of Schaffer collaterals; orange: OLM cells; red: basket cells; blue: pyramidal cells). The last panels display spectrograms of the firing rate of each population throughout the simulation.

### Single cell response to extracellular stimulation

After setting up the model and tuning its parameters such as it exhibits theta-nested gamma oscillations, we sought to study the effects of extracellular electrical stimulation, first on single-cell activity and then on network dynamics. We first aimed to confirm the influence of electrode location and stimulation polarity on the different cell types individually (Fig. 4). To this end, for each cell type, we performed several 350ms simulations during which a 1ms single-pulse stimulation was applied to the extracellular space surrounding an individual neuron. The electrode was positioned at various locations in the x-y plane, with an increment of 50 µm in both x and y directions. We tested different stimulation amplitudes, ranging from 0.01 mA to 10 mA in steps of 0.01 mA, and recorded the smallest amplitude that elicited a somatic action potential (stimulation amplitude threshold).

**Fig 4.**
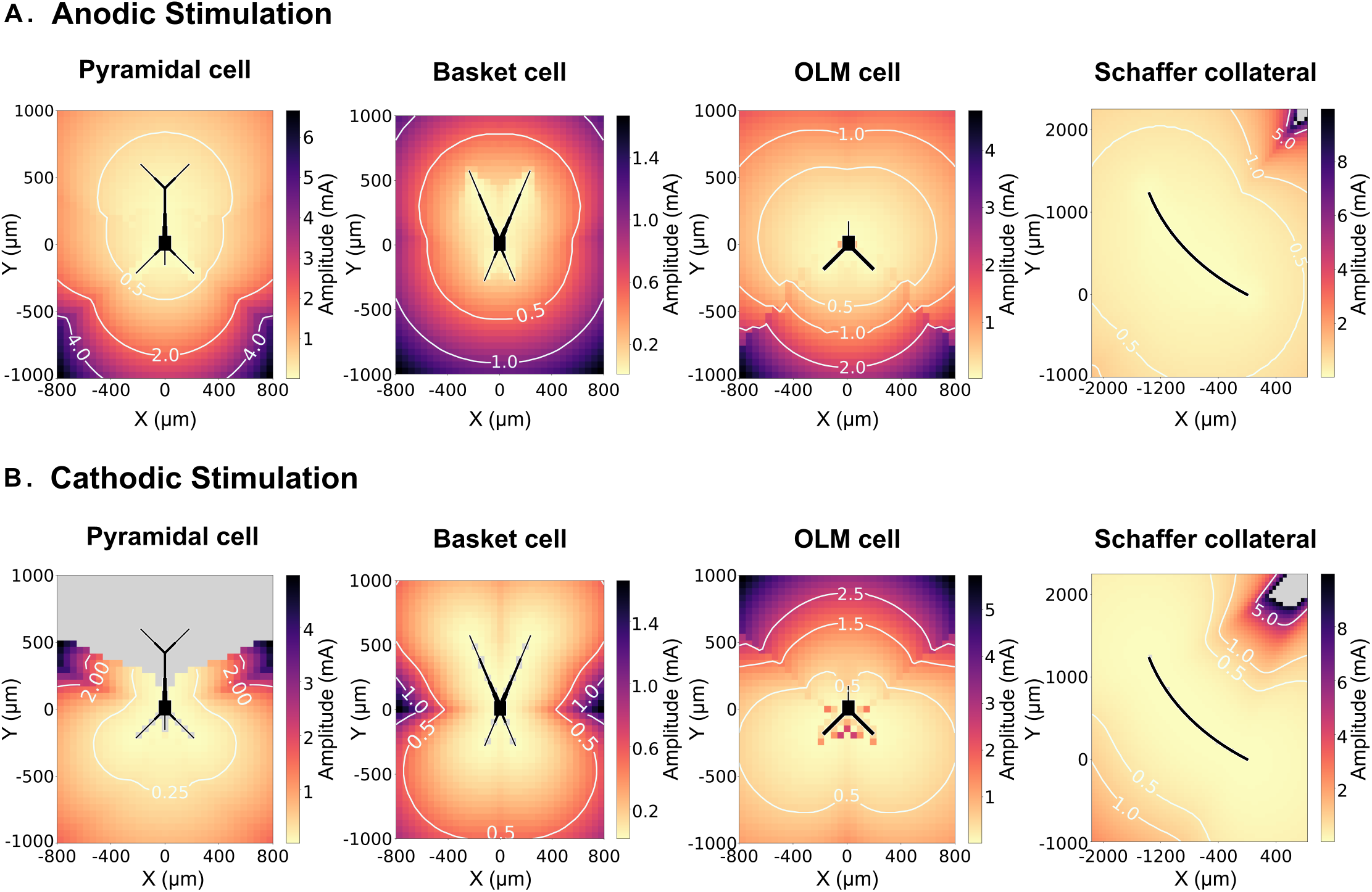
Stimulation threshold of single cell. A 1-ms single-pulse monopolar stimulation was applied to the extracellular space surrounding individual neuron. Several simulations were performed with the stimulation amplitude increased from 0.01 to 10 mA in steps of 0.01 mA. The electrode was positioned at various locations in the x-y plane with a resolution of 50 µm (i.e. electrode positions were sampled every 50 µm in both x and y directions). For each electrode position, we recorded the smallest amplitude that elicited a somatic action potential. **(A)** Stimulation amplitude threshold as a function of neuron-electrode distance for anodic stimulation for each cell type. **(B)** Stimulation amplitude threshold as a function of neuron-electrode distance for cathodic stimulation for each cell type. The gray areas indicate electrode positions for which no action potential could be elicited, either because the required stimulation amplitude exceeded 10 mA, or because the neuron was too sensitive. The results showed that the stimulation amplitude threshold depended on the stimulation polarity. Pyramidal and OLM cells exhibited inverted response profiles between cathodic and anodic stimulation. Basket cells showed greater sensitivity to anodic stimulation compared to cathodic stimulation. Overall, Schaffer collaterals were the most responsive to extracellular stimulation, followed by basket cells.

#### Electrode location

The results presented in Fig. 4 demonstrate that the stimulation amplitude threshold increases with the electrode-to-neuron distance for both anodic and cathodic stimulation across all cell types. For example, in the case of anodic stimulation applied to the extracellular space of a pyramidal neuron (first panel of Fig. 4A), the current amplitude needed to elicit a spike in the soma of the neuron increased when the electrode was moved further away from the neuron.

Furthermore, the morphology of the neuron influences the stimulation threshold. The increase in threshold with the distance from the neuron was not uniform across its surface. For anodic stimulation, the threshold rose more sharply around the basal dendrites of the pyramidal neuron than around its apical dendrites. The same phenomenon was observed for the OLM cell (third panel of Fig. 4A): the stimulation amplitude threshold rose faster along the dendrites and more smoothly around the axon.

We also showed that the stimulation threshold depends on the cell type because some cells were more excitable than others. In the case of anodic stimulation (Fig. 4A), at the same electrode position, a lower current amplitude was required to elicit an action potential in the basket cell than in pyramidal and OLM cells. The stimulation threshold of the Schaffer collateral was lower than those of the three cell types for both cathodic and anodic stimulation.

#### Stimulation polarity

Stimulation threshold profiles varied with stimulation polarity across all cell types. In basket cells, the stimulation threshold increased more gradually with distance from the neuron under cathodic stimulation than under anodic stimulation, consistent with the greater excitatory effect of extracellular cathodic stimulation compared to anodic stimulation [49]. For OLM cells, anodic stimulation elicited action potentials at lower current amplitudes near the axonal projection than near the dendritic arbor. In contrast, under cathodic stimulation, lower current amplitudes were required near the dendritic arbor, whereas higher amplitudes were needed toward the axonal projection.

### Population response to extracellular stimulation

Our main objective was then to study the influence of various stimulation parameters on the network oscillatory activity.

#### Stimulation amplitude, electrode position and orientation

We first analyzed the effects of stimulation amplitude and electrode configuration (Figs. 5, 6). To this end, we performed several 5-second simulations in which a 2-second 50 Hz biphasic extracellular stimulation pulse train was applied to the network. We tested different electrode configurations, including monopolar and bipolar arrangements. For each configuration, we explored a wide range of stimulation amplitudes, from 0.5 mA to 6 mA in 0.5 mA increments, and quantified for each neuronal type the proportion of neurons that significantly increased or decreased their firing rate during the stimulation period compared to their baseline activity. The simulation results showed that the network activity increased with the stimulation amplitude across all stimulation arrangements. We also observed that for both monopolar (Fig. 5) and bipolar (Fig. 6) configurations, stimulation applied to the inner curvature of the hippocampal CA1 region induced stronger network excitation than stimulation on the outer curvature. For instance, in the case of monopolar stimulation at 3 mA, around 65% of pyramidal cells increased their activity when the electrode was placed on the outer curvature of CA1 (Fig. 5B1) against ∼90% when the electrode was placed on the inner curvature (Fig. 5B2). For a stimulation amplitude of 6 mA, ∼85% of pyramidal cells increased their activity when the electrode was placed on the outer curvature, while the percentage reached 100% when placed on the inner curvature. This difference in recruitment depending on electrode placement was more pronounced for bipolar orthogonal stimulation with at most ∼65% of pyramidal cells increasing their activity when the electrode was positioned on the outer curvature (Fig. 6A1), against 100% when positioned on the inner curvature (Fig. 6A3).

**Fig 5.**
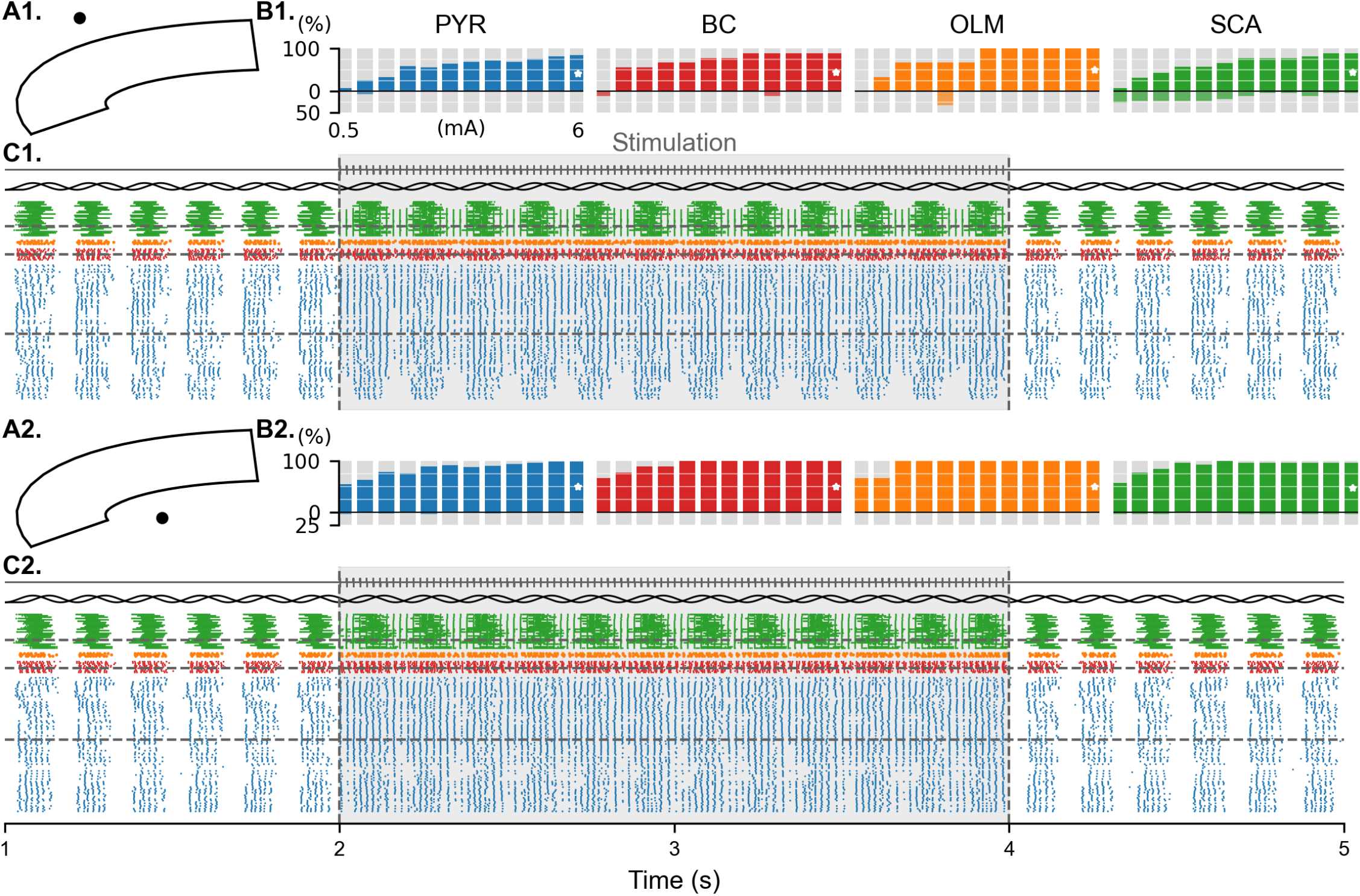
Network response to variation of stimulation amplitude with monopolar electrodes. **(A)** Spatial positioning of the stimulating electrode with respect to the CA1 slice. The electrode is depicted as a black dot indicating monopolar stimulation. **(B)** Percentage of neurons in each population exhibiting increased or decreased spiking activity during stimulation compared to baseline. The x-axis shows stimulation amplitude ranging from 0.5 to 6 mA in 0.5 mA steps. The y-axis represents the percentage of neurons showing a change in activity. The horizontal black line at 0% denotes baseline activity: upward shifts indicate percentage of neurons showing an increased activity, while downward shifts indicate the percentage of neurons showing a decreased activity. **(C)** Simulation results obtained at a stimulation amplitude of 6 mA, corresponding to the white stars in panel B. The top part displays the stimulation waveform which is a biphasic monopolar stimulation at 50 Hz. The pulse-width is set to 300 µs with an interphase of 100 µs. The vertical dashed lines mark the beginning (at 2 seconds of simulation) and end (at 4 seconds) of stimulation. The black sinusoidal traces represent the theta-frequency oscillatory inputs injected into the Schaffer collaterals. The bottom part shows neuronal spiking activity over time for each population (green: last nodes of Schaffer collaterals; orange: OLM cells; red: basket cells; blue: pyramidal cells). Horizontal dashed gray lines indicate the neuron closest to the stimulating electrode in each population. The results revealed a greater increase in activity when the electrode was placed within the inner curvature of the CA1 slice.

**Fig 6.**
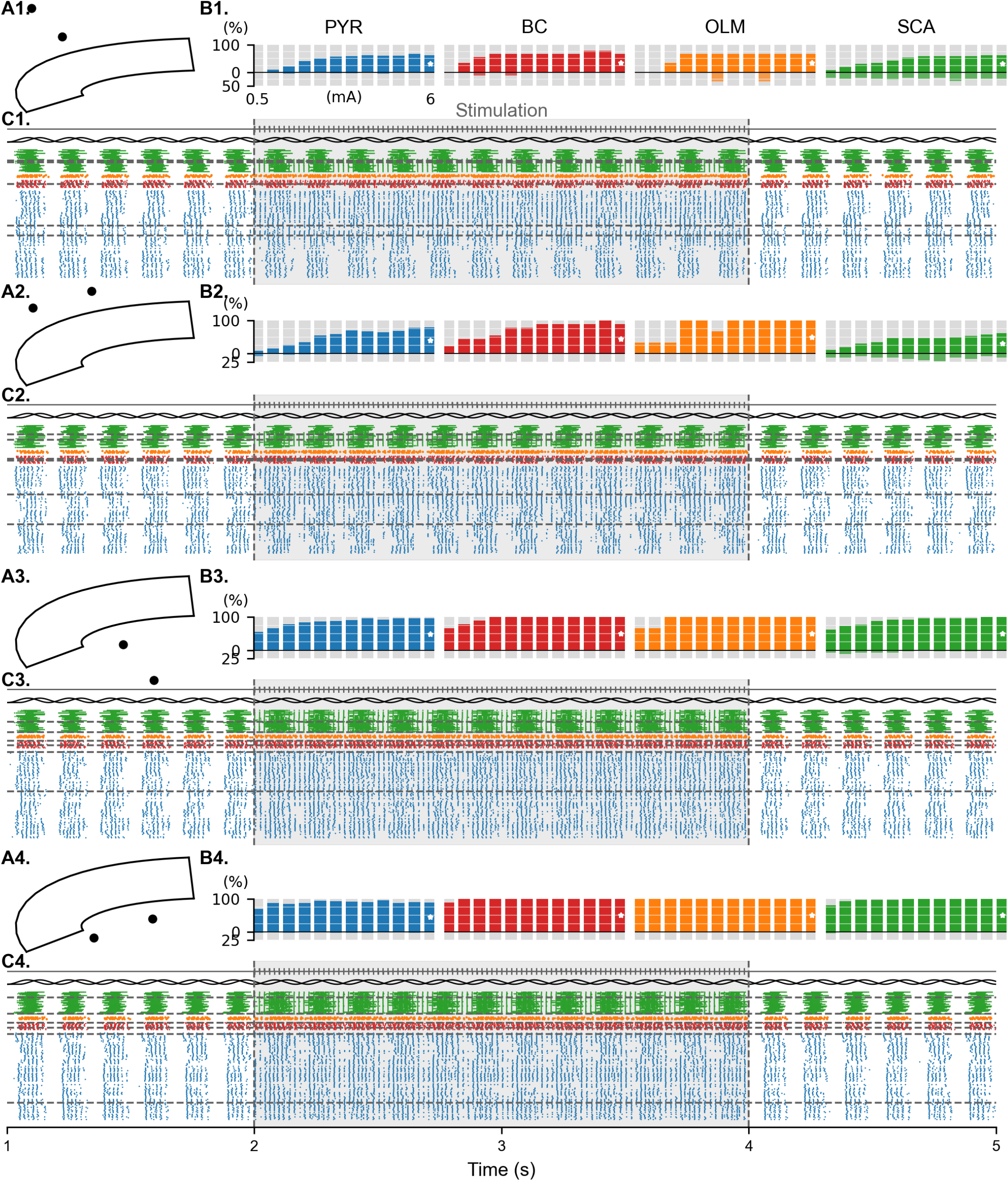
Network response to variation of stimulation amplitude with bipolar electrodes. **(A)** Spatial positioning of the stimulating electrodes with respect to the CA1 slice. The electrode are depicted as two black dots separated by 1.5 mm, indicating bipolar stimulation. **(B)** Percentage of neurons in each population exhibiting increased or decreased spiking activity during stimulation compared to baseline. The x-axis shows stimulation amplitude ranging from 0.5 to 6 mA in 0.5 mA steps. The y-axis represents the percentage of neurons showing a change in activity. The horizontal black line at 0% denotes baseline activity: upward shifts indicate percentage of neurons showing an increased activity, while downward shifts indicate the percentage of neurons showing a decreased activity. **(C)** Simulation results obtained at a stimulation amplitude of 6 mA, corresponding to the white stars in panel B. The top part displays the stimulation waveform which is a biphasic monopolar stimulation at 50 Hz. The pulse-width is set to 300 µs with an interphase of 100 µs. The vertical dashed lines mark the beginning and end of stimulation. The black sinusoidal traces represent the theta-frequency oscillatory inputs injected into the Schaffer collaterals. The bottom part shows neuronal spiking activity over time for each population (green: last nodes of Schaffer collaterals; orange: OLM cells; red: basket cells; blue: pyramidal cells). Horizontal dashed gray lines indicate the neuron closest to the stimulating electrode in each population. The results show that the neuronal recruitment was influenced by both electrode placement and orientation. Positioning the electrodes within the inner curvature of CA1 evoked a more pronounced increase in network activity than positioning the electrodes on the outer curvature. Placing the electrodes parallel to the Schaffer collaterals (panels A2 and A4) induced a greater increase in activity than placing them perpendicular to the Schaffer collaterals (panels A1 and A3).

Furthermore, in the case of bipolar stimulation, we found that placing the electrodes parallel to the Schaffer collaterals (Fig. 6A2 and A4) induced a greater increase in network activity than when the electrodes were oriented orthogonal to the Schaffer collaterals (Fig. 6A1 and A3). For a stimulation amplitude of 6 mA, ∼80% of pyramidal cells increased their spiking activity for parallel electrode orientation (Fig. 6B2), against ∼60% for orthogonal electrode orientation (Fig. 6B1).

With bipolar electrodes placed on the outer curvature of the hippocampus (and with the orthogonal montage in particular), it appeared that even with high stimulation amplitude, some pyramidal and Schaffer collaterals were not increasing their firing rate during the stimulation period. Indeed, even with a stimulation amplitude of 6 mA, only ∼60% of pyramidal and Schaffer collaterals increased their spiking activity (Fig. 6B1). Instead, in this configuration, 25% of the Schaffer collaterals reduced their spiking activity during the stimulation period.

#### Stimulation frequency and shape

Next, we investigated the effect of the stimulation frequency on the network response. The electrodes were placed on the outer curvature of the hippocampal CA1, parallel to the Schaffer collaterals (see Fig. 7D), with the stimulation amplitude fixed at 3.5 mA. We compared bipolar biphasic pulse trains at 50 Hz and 130 Hz against theta-burst stimulation at 5 Hz. The results show that around 75% of pyramidal cells increased their activity for 50-Hz and 130-Hz stimulation, while the percentage only reached 25% for theta-burst stimulation. However, the effects induced by theta-burst stimulation seemed to depend on stimulation timing relative to the ongoing theta.

**Fig 7.**
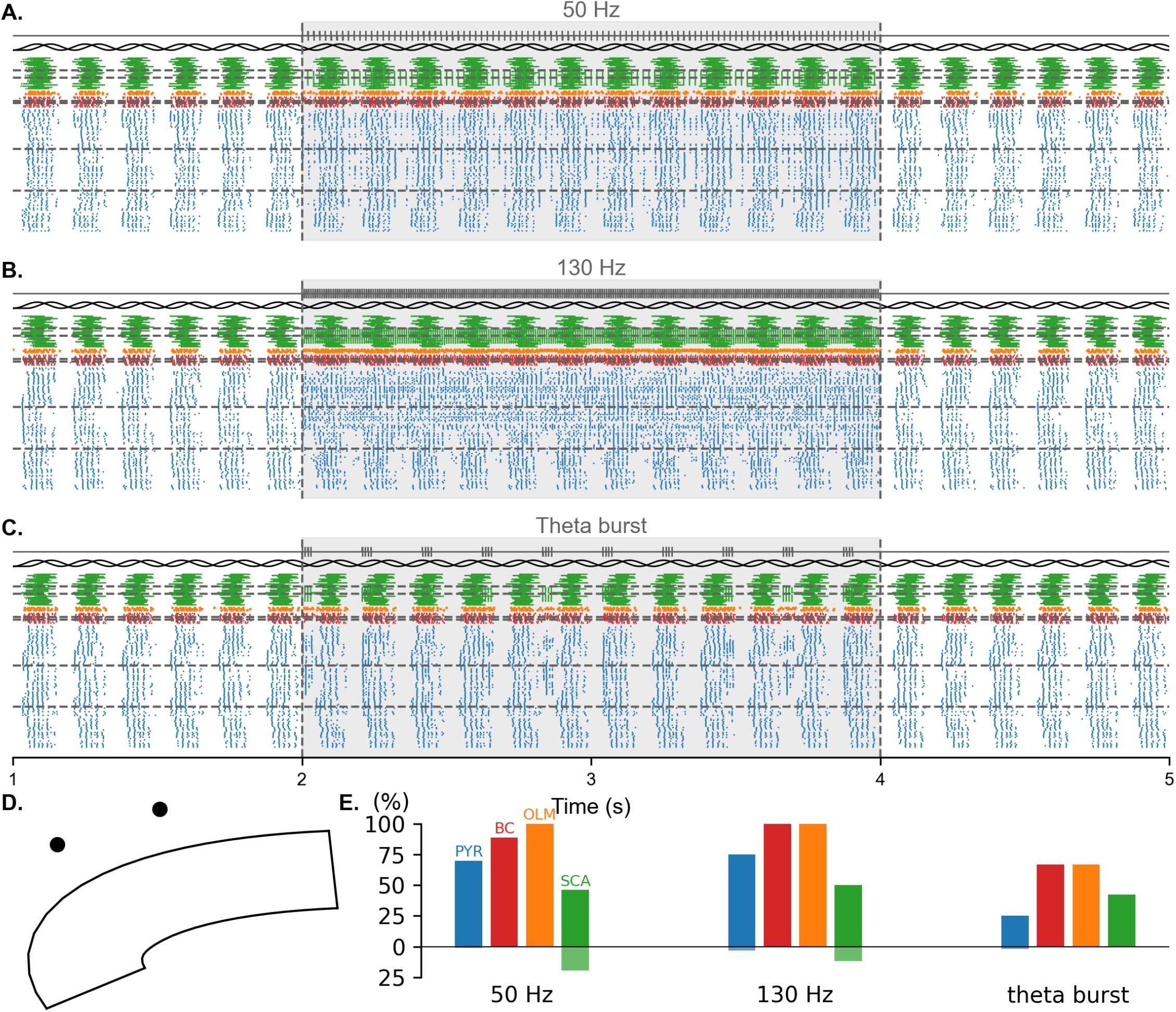
Network response to variation of frequency. All simulations results were obtained with a stimulation amplitude of 3.5 mA. **(A)** Neuronal spiking activity in the network in response to a bipolar biphasic stimulation at 50 Hz. The top part displays the stimulation waveform. The vertical dashed lines mark the beginning and end of stimulation. The black sinusoidal traces represent the theta-frequency oscillatory inputs injected into the Schaffer collaterals. The bottom part shows neuronal spiking activity over time for each population (green: last nodes of Schaffer collaterals; orange: OLM cells; red: basket cells; blue: pyramidal cells). Horizontal dashed gray lines indicate the neuron closest to the stimulating electrode in each population. **(B)** Neuronal spiking activity in the network in response to a bipolar biphasic stimulation at 130 Hz. **(C)** Neuronal spiking activity in the network in response to a bipolar biphasic theta-burst stimulation. The pulse-width is set to 200 µs with an interphase of 100 µs. 4 biphasic pulses at a frequency of 100 Hz were presented every 200 ms. **(D)** Spatial positioning of the stimulating electrodes with respect to the CA1 slice. **(E)** Percentage of neurons in each population exhibiting increased or decreased spiking activity during stimulation, depending on the stimulation parameters. The y-axis represents the percentage of neurons showing a change in activity. The horizontal black line at 0% denotes baseline activity: upward shifts indicate percentage of neurons showing an increased activity, while downward shifts indicate the percentage of neurons showing a decreased activity. The results reveal that 130 Hz stimulation induced a greater increase in network activity compared to 50 Hz stimulation or theta-burst stimulation. Theta-burst elicited a less pronounced global increase in activity but the stimulation effects appeared to depend on the timing of stimulation relative to the phase of ongoing theta oscillations.

#### Presence or absence of Schaffer collaterals

Finally, while stimulation in the presence of Schaffer collaterals resulted in increased activity in all cell types, their absence elicited mixed excitatory and inhibitory responses (Fig. 8). The inhibitory effect on pyramidal cells was more pronounced at 130 Hz, with 25% of pyramidal neurons showing a decrease in activity, than at 50 Hz where only ∼10% of pyramidal cells had a decrease in activity. These findings suggest that the net increase in network activity observed during stimulation is primarily driven by excitatory inputs from Schaffer collaterals. In the absence of Schaffer collaterals, basket cells are the most sensitive to extracellular stimulation and could therefore induce inhibition of the pyramidal cells.

**Fig 8.**
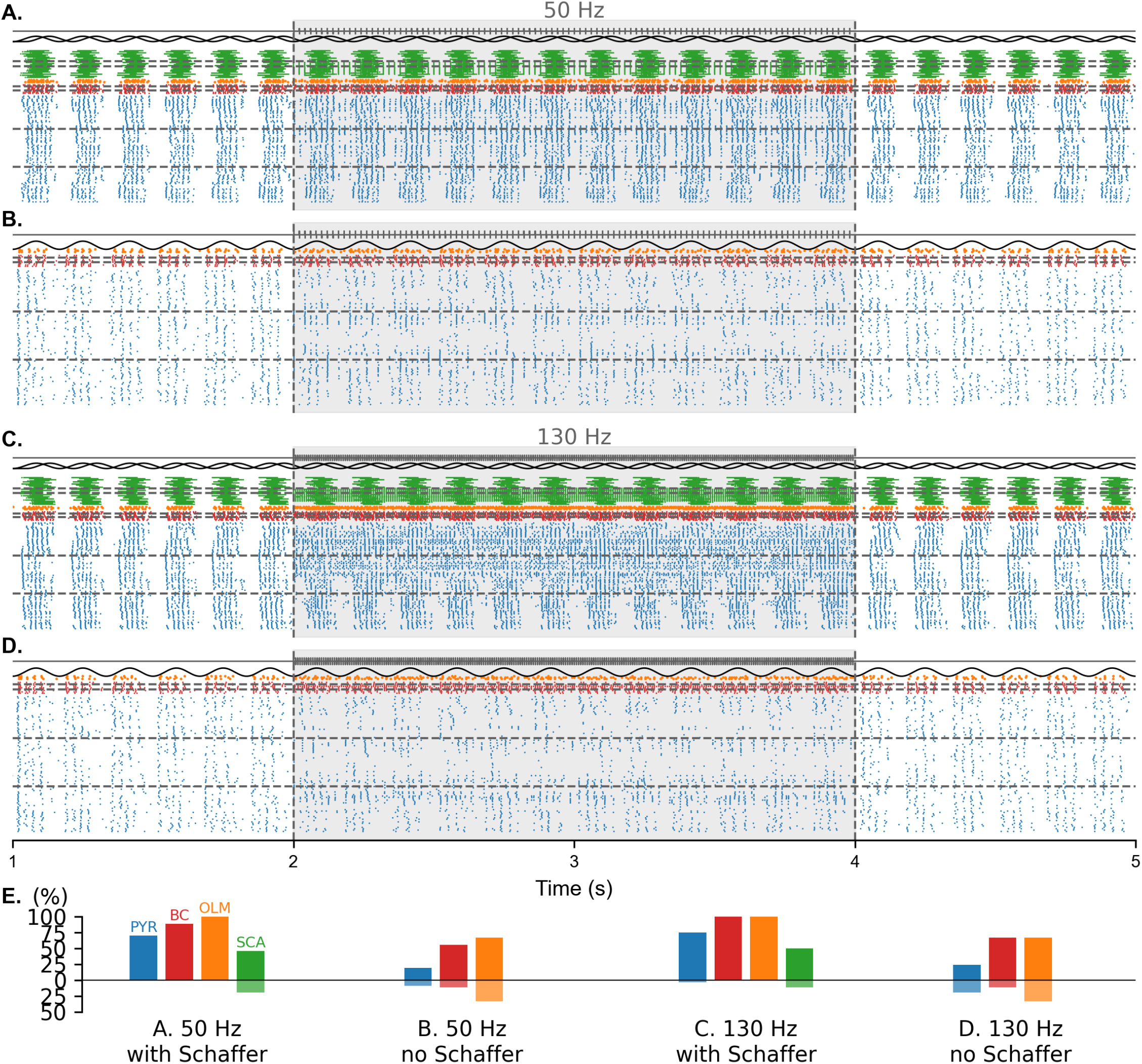
Network response to electrical stimulation with and without Schaffer collaterals. All simulations results were obtained with a stimulation amplitude of 3.5 mA. Stimulating electrodes were bipolar and placed along the outer curvature of CA1, parallel to the Schaffer collaterals as in Figure 7D. **(A)** Spiking activity of the neurons in the network when the pyramidal cells receive inputs from the Schaffer collaterals. The top part displays the stimulation waveform, which was a 50 Hz bipolar biphasic pulse train. The vertical dashed lines mark the beginning and end of stimulation. The black sinusoidal traces represent the theta-frequency oscillatory inputs injected into the Schaffer collaterals. The bottom part shows neuronal spiking activity over time for each population (green: last nodes of Schaffer collaterals; orange: OLM cells; red: basket cells; blue: pyramidal cells). Horizontal dashed gray lines indicate the neuron closest to the stimulating electrode in each population. **(B)** Spiking activity of the neurons in the network when Schaffer collaterals are removed from the network. Stimulation parameters are the same as in panel A. Theta-oscillatory inputs are injected directly into the pyramidal cells. **(C)** Simulation results obtained with a stimulation frequency of 130 Hz in the full network model, including the Schaffer collaterals. **(D)** Simulation results obtained with a stimulation frequency of 130 Hz when the Schaffer collaterals are removed from the network. **(E)** Percentage of neurons in each population exhibiting increased or decreased spiking activity during stimulation, depending on the stimulation parameters. The y-axis represents the percentage of neurons showing a change in activity. The horizontal black line at 0% denotes baseline activity: upward shifts indicate percentage of neurons showing an increased activity, while downward shifts indicate the percentage of neurons showing a decreased activity. The results showed that 130 Hz stimulation induced a greater increase in network activity compared to 50 Hz stimulation, both in networks with and without inputs from Schaffer collaterals. Furthermore, in the absence of Schaffer collateral input, pyramidal cells activity decreased more prominently, particularly under 130 Hz stimulation.

## Discussion

### Highlights

In this work, we have developed a reduced multicompartment network model of CA1 that exhibits theta-nested gamma oscillations. Our model incorporates multicompartment neurons and explicit axonal afferents to meaningfully simulate the effects of extracellular stimulation of the CA1 area [33, 34, 50, 51]. Simplifications in the neuronal anatomy, as well as the number of neurons and connectivity, enable computationally tractable simulations on a portable computer. This balance between accuracy and efficiency allows for the systematic testing of various stimulation configurations within a reasonable timeframe. We believe this work provides a template that can be used to gain intuition regarding the effects of other stimulation configurations, the effects of stimulation on pathologies, or even a foundation for building customized reduced models.

In order to build a CA1 model that generates theta-gamma oscillations, we initially applied the scaling method described by Romaro and colleagues [52] to downscale the network implemented by Vardalakis and colleagues [32]. However, this approach failed to reproduce gamma oscillations. This is likely due to the increase in neuronal morphological complexity, which probably altered the intrinsic excitability of the cells. Furthermore, our reduced network size may have reached the limits of the scaling methodology used by Romaro and colleagues. While they suggest compensating the loss of synaptic input by direct current, we found this approach computationally prohibitive. Instead, we iteratively adjusted the synaptic weights in the CA1 network in the absence of Schaffer collaterals until gamma oscillations emerged (see Fig. 2). Then, we added the Schaffer collaterals to the network in order to introduce theta input. We optimized both theta amplitude and the synaptic weight from the collaterals to the CA1 network in order to get strong theta-gamma oscillations (see Fig. 3).

We investigated the response of individual cells to extracellular stimulation. Our results demonstrate that for all cell types, the stimulation amplitude threshold increases with electrode-to-neuron distance, regardless of stimulation polarity. Moreover, neuronal morphology modulates these thresholds, as the amplitude threshold depends on the location of the electrode alongside the different neuronal elements. The stimulation polarity also influences the threshold profiles. All these results are in line with previous computational work of McIntyre and Grill [53]. Additionally, our simulations validate that axonal fibers exhibit higher sensitivity to extracellular stimulation than cell bodies [34]. As shown in Figure 4, Schaffer collaterals exhibit smaller thresholds compared to other cell types. Given the scarcity of studies analyzing cell-type-specific responses to extracellular stimulation, our study provides predictions, particularly for the interneuron populations. However, because most neuron models are calibrated for intracellular stimulation, their predicted response to extracellular stimulation should be interpreted with caution. Finally, the grey regions in the first panel of Figure 4B denote electrode locations where action potentials failed to initiate despite increasing the stimulation amplitude. The observed effect may result from numerical instability arising from a dual contribution of electrode proximity and the use of explicit Euler integration in certain equations.

We also examined the network response to stimulation by comparing conditions with and without Schaffer collateral inputs (Fig. 8). Our results demonstrate that the presence of Schaffer collaterals enhances the activity of pyramidal cells, whereas their removal attenuates their response. This observation supports the hypothesis that DBS acts indirectly on local neurons by recruiting afferent pathways [51, 54, 55]. Specifically, the excitation of Schaffer collaterals in response to DBS serves to drive the activity of the underlying pyramidal cell population. In the absence of Schaffer collaterals, DBS acts directly on the targeted brain area. It has been hypothesized that DBS exerts direct inhibition on local cells, either by depolarization block, or by activating GABAergic pathways [54, 56]. In our work, GABAergic afferent pathways were not represented. Therefore, the decrease of activity in pyramidal neurons could originate from depolarization block or from the direct recruitment of local interneurons which subsequently inhibit the pyramidal population.

### Limitations

Since our model serves as an initial template for building computationally efficient yet accurate models, only the most crucial mechanisms involved in theta-nested gamma oscillations and extracellular stimulation have been implemented. Therefore, many assumptions and simplifications had to be made, and further improvements could be implemented. For instance, theta oscillations are provided as input to the network.

Numerous studies have suggested that theta oscillations in the CA1 area of the hippocampus originate from inputs from neighboring regions [31, 57–59]. Specifically, it is thought that theta emerges from the interplay between inputs from CA3 Schaffer collaterals, entorhinal cortex perforant path and medial septum GABAergic projections. Although theta oscillations in CA1 can be entrained by extrinsic inputs, local cells can also be responsible for generating theta. For example, Colgin [58] suggests that theta can emerge from local interactions between pyramidal and inhibitory neurons. Some computational studies have also investigated the emergence of theta rhythms within CA1 and found that theta can be generated endogenously within the network [27, 28]. Bezaire and colleagues [28] have suggested that the interconnection between several pyramidal neurons is sufficient to generate theta oscillations within the network.

However, given the scale of our network and the fact that we did not account for pyramidal-to-pyramidal connections, this theta-generation mechanism could not be implemented in our model. Furthermore, Sengupta and colleagues [27] have proposed that theta emerges from the interactions between pyramidal and bistratified cells. For simplification purposes, we did not implement all known interneuron types in CA1, such as bistratified cells. Therefore, we could not test this theory either.

Another limitation is the exclusion of CA1 input pathways besides the Schaffer collaterals. CA1 pyramidal cells also receive excitatory inputs from the entorhinal cortex through the perforant path. The medial septum also provides inhibitory inputs to CA1. Since the effects of DBS primarily depend on the activated pathways [55], modeling projections from both the entorhinal cortex and the medial septum could yield a more complex and accurate response to stimulation. Furthermore, as these different pathways do not innervate the same layers of CA1, electrode placement and orientation could have a more significant impact on the network’s response to stimulation. In addition, the Schaffer collaterals in our model maintain a constant diameter and do not include branching points. However, it is well-established that hippocampal axonal projections taper and branch. These features significantly impact how they are recruited during stimulation [36]. While incorporating these morphological characteristics would increase the computational cost of our model, doing so would be essential for investigating the recruitment of the different pathways under varying electrode and stimulation configurations.

In addition, while this study focused on the acute effects of DBS, we did not account for activity-dependent plasticity mechanisms operating at different timescales. At short timescales, stimulation of afferent fibers triggers action potentials that propagate to the axonal terminals where neurotransmitters are released into the synaptic cleft. However, during high-frequency DBS, the rate at which action potentials are generated outpaces the biological mechanisms responsible for vesicle replenishment. This leads to synaptic depression. Consequently, the postsynaptic neurons fail to reach threshold, attenuating signal transmission [51, 60, 61]. Incorporating these short-time plasticity mechanisms could substantially alter how the network responds to sustained high-frequency stimulation. Furthermore, chronic DBS is known to induce persistent changes in neural circuits through long-term plasticity mechanisms. In particular, repeated stimulation may reshape connectivity through spike-timing-dependent plasticity. Integrating such mechanisms is essential to capturing how stimulation affects the network beyond the stimulation period and for investigating whether DBS can induce durable network reorganization. Finally, while this study tested the effects of stimulation in a healthy network, it would also be interesting to investigate whether stimulation can restore theta-gamma PAC in pathological networks and whether plasticity mechanisms contribute to the recovery and maintenance of physiological oscillatory dynamics.

### Perspectives

Future work could focus on expanding the model to encompass the entire hippocampal formation while explicitly modeling the different pathways. Such an expansion would allow for the investigation of various stimulation targets and their effects on downstream hippocampal regions, as well as on global theta-gamma oscillations within the hippocampus. Given that closed-loop stimulation represents a highly promising approach, future studies should focus on implementing these protocols to evaluate their impact on oscillatory dynamics. Furthermore, by introducing pathological state in the network, we could determine the optimal stimulation strategies for restoring theta-gamma coupling. Another essential addition would be to incorporate mechanisms of synaptic plasticity in order to evaluate the effect of extracellular stimulation after the end of the stimulus.

## Materials and methods

### Neuron models

Our model is composed of pyramidal neurons, PV+ basket cells and OLM cells. All cells follow the Hodgkin-Huxley formalism [62] and were replicated from previous work of Bezaire and colleagues [28]. Each cell type is modeled with multiple compartments (so as to be able to account for extracellular stimulation) but with reduced morphology (see Fig. 1A) to balance simulations’ accuracy and computational cost. We also represented the Schaffer collaterals projecting from the CA3 area of the hippocampus to the CA1 area as myelinated axons, using a multicompartment double cable model following the Hodgkin-Huxley formalism and replicated from the previous work of McIntyre and colleagues [38]. All neurons and collaterals were implemented using NEURON 8.2 with Python 3.10.9 [63]. For each compartment, the evolution of the membrane potential *V*_*m*_ is governed by:

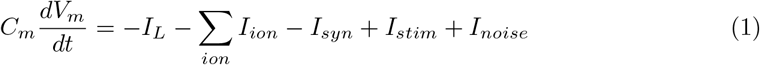

where *C*_*m*_ is the membrane capacitance, *I*_*L*_ is the leak current, *I*_*ion*_ represents the ionic currents associated with specific ion channels present in the compartment, *I*_*syn*_ denotes the synaptic input, *I*_*stim*_ is the stimulus current applied to the compartment and *I*_*noise*_ is a stochastic current representing background fluctuations.

*I*_*noise*_ is injected into all compartments of all cells, except axon of pyramidal cells. Schaffer collaterals are also excluded from noise injection. The added noise is a Ornstein-Uhlenbeck process described by the following equation:

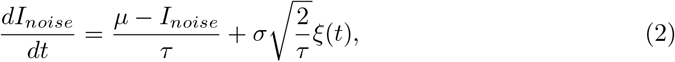

where *µ* is the mean value, *τ* is the time constant governing the rate at which the process reverts to the mean, *σ* is the noise amplitude, and *ξ*(*t*) represents Gaussian white noise with 0 mean and unit variance. We tested various noise amplitudes for each cell type and selected values that elicited minimal spontaneous activity. This procedure resulted in a noise amplitude of 0.01 nA for pyramidal and basket cells, and a lower amplitude of 0.0025 nA for the more sensitive OLM cells.

The leak current follows the equation:

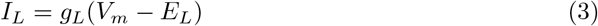

where *g*_*L*_ is the maximum conductance of the leak channel and *E*_*L*_ corresponds to its reversal potential.

All ionic current equations for each cell type were taken from the model of Bezaire and colleagues [28] and are listed below. Although all equations are written in a continuous form here, some were implemented using an explicit Euler integration method, as in [28].

#### Pyramidal cells

Pyramidal cells are modeled with 83 compartments organized into somatic, basal, apical and axonal groups. All compartments contain a sodium current (*I*_*Na*_), a delayed rectifier potassium current (*I*_*K,dr*_) and an A-type potassium current (*I*_*K,A*_). Apical dendrites include an additional A-type current (*I*_*K,Adist*_) which applies to dendrites at least 100-µm away from the soma, while *I*_*K,A,apical*_ applies only to the dendrites that are within 100 µm of the soma. Somatic and apical compartments also contain a hyperpolarization-activated cyclic nucleotide-gated current (*I*_*HCN*_).

The sodium current *I*_*Na*_ is described by:

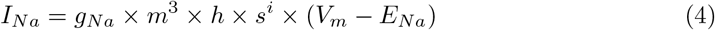

where *g*_*Na*_ is the maximum conductance, *E*_*Na*_ is the reversal potential and *m, h* and *s* are gating variables defined by the equations:

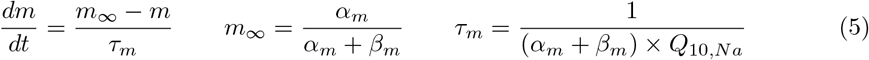

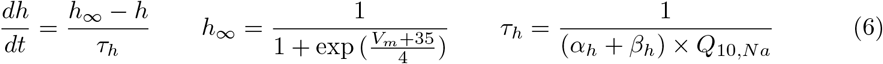

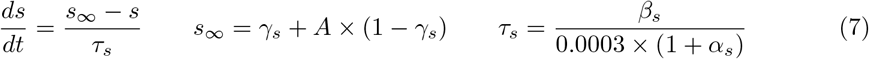

The potassium currents are described by:

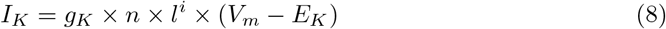

where *g*_*K*_ is the maximum conductance, *E*_*K*_ is the reversal potential, *n* and *l* are gating variables defined by the equations:

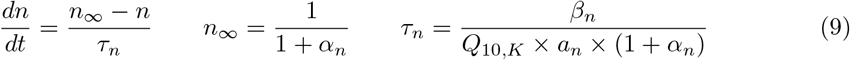

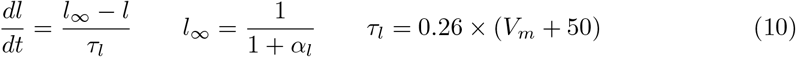

*I*_*HCN*_ current is described by:

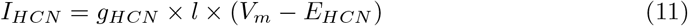

where *g*_*HCN*_ is the maximum conductance, *E*_*HCN*_ is the reversal potential and *l* is the gating variable defined by:

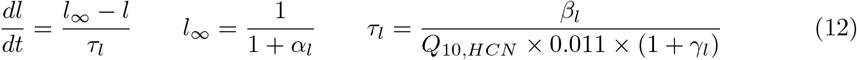

The full expressions for the parameters described above can be found in Table 1.

**Table 1.** Parameter expressions for pyramidal neurons.

| Parameter | Expression |
| --- | --- |
| $C_{m,soma,axon}$ | 1 $\mu F/cm^2$ |
| $C_{m,basal,apical}$ | 2 $\mu F/cm^2$ |
| $C^\circ$ | 35°C |
| $g_{L,soma,axon}$ | 0.00004 $S/cm^2$ |
| $g_{L,basal,apical}$ | 0.00007 $S/cm^2$ |
| $E_L$ | -66 $mV$ |
| $g_{Na,soma,basal,apical}$ | 0.032 $S/cm^2$ |
| $g_{Na,axon}$ | 0.064 $S/cm^2$ |
| $E_{Na}$ | 55 $mV$ |
| $i_{Na,soma,basal,apical}$ | 1 |
| $i_{Na,axon}$ | 0 |

Table 1. Parameter expressions for pyramidal neurons (continued)
| Parameter | Expression |
| --- | --- |
| $A_{Na,soma,basal}$ | 1 |
| $A_{Na,apical}$ | 0.8 |
| $\alpha_{m,Na}$ | $\frac{0.4 \times (V_m + 15)}{1 - \exp\left(\frac{-(V_m + 15)}{7.2}\right)}$ |
| $\beta_{m,Na}$ | $\frac{-0.124 \times (V_m + 15)}{1 - \exp\left(\frac{(V_m + 15)}{7.2}\right)}$ |
| $\alpha_{h,Na}$ | $\frac{0.03 \times (V_m + 30)}{1 - \exp\left(\frac{-(V_m + 30)}{1.5}\right)}$ |
| $\beta_{h,Na}$ | $\frac{-0.01 \times (V_m + 30)}{1 - \exp\left(\frac{(V_m + 30)}{1.5}\right)}$ |
| $\alpha_{s,Na}$ | $\exp\left(\frac{1.0e^{-3} \times 12 \times (V_m + 45) \times 9.648e^4}{8.315 \times (273.16 + C^\circ)}\right)$ |
| $\beta_{s,Na}$ | $\exp\left(\frac{1.0e^{-3} \times 2.4 \times (V_m + 45) \times 9.648e^4}{8.315 \times (273.16 + C^\circ)}\right)$ |
| $\gamma_{s,Na}$ | $\frac{1}{1 + \exp\left(\frac{V_m + 43}{2}\right)}$ |
| $Q_{10,Na}$ | $2^{\frac{C^\circ - 24}{10}}$ |
| $g_{K,dr}$ | 0.003 $S/cm^2$ |
| $E_{K,dr}$ | -90 $mV$ |
| $i_{K,dr}$ | 0 |
| $\alpha_{n,K,dr}$ | $\exp\left(\frac{-1.0e^{-3} \times 3 \times (V_m - 13) \times 9.648e^4}{8.315 \times (273.16 + C^\circ)}\right)$ |
| $\beta_{n,K,dr}$ | $\exp\left(\frac{-1.0e^{-3} \times 2.1 \times (V_m - 13) \times 9.648e^4}{8.315 \times (273.16 + C^\circ)}\right)$ |
| $Q_{10,K,dr}$ | $1^{\frac{C^\circ - 24}{10}}$ |
| $a_{n,K,dr}$ | 0.02 |
| $g_{K,A,soma,basal}$ | 0.008 $S/cm^2$ |
| $g_{K,A,axon}$ | 0.0016 $S/cm^2$ |
| $g_{K,A,apical}$ | $0.008 \times \left(\frac{1 + dist_{soma}}{100}\right)$ $S/cm^2$ |
| $E_{K,A}$ | -90 $mV$ |
| $i_{K,A}$ | 1 |
| $\alpha_{n,K,A}$ | $\exp\left(\frac{1.0e^{-3} \times \zeta \times (V_m - 11) \times 9.648e^4}{8.315 \times (273.16 + C^\circ)}\right)$ |

Table 1. Parameter expressions for pyramidal neurons (continued)
| Parameter | Expression |
| --- | --- |
| $\beta_{n,K,A}$ | $\exp\left(\frac{1.0e^{-3} \times \zeta \times 0.55 \times (V_m - 11) \times 9.648e^4}{8.315 \times (273.16 + C^\circ)}\right)$ |
| $\zeta$ | $1.5 + \frac{-1}{1 + \exp\left(\frac{V_m + 40}{5}\right)}$ |
| $\alpha_{l,K,A}$ | $\exp\left(\frac{1.0e^{-3} \times 3 \times (V_m + 56) \times 9.648e^4}{8.315 \times (273.16 + C^\circ)}\right)$ |
| $Q_{10,K,A}$ | $5^{\frac{C^\circ - 24}{10}}$ |
| $a_{n,K,A}$ | 0.05 |
| $g_{K,Adist}$ | $0.008 \times \left(\frac{1 + dist_{soma}}{100}\right) \quad S/cm^2$ |
| $E_{K,Adist}$ | -90 $mV$ |
| $i_{K,Adist}$ | 1 |
| $\alpha_{n,K,Adist}$ | $\exp\left(\frac{1.0e^{-3} \times \zeta \times (V_m + 1) \times 9.648e^4}{8.315 \times (273.16 + C^\circ)}\right)$ |
| $\beta_{n,K,Adist}$ | $\exp\left(\frac{1.0e^{-3} \times \zeta \times 0.39 \times (V_m + 1) \times 9.648e^4}{8.315 \times (273.16 + C^\circ)}\right)$ |
| $\zeta$ | $1.5 + \frac{-1}{1 + \exp\left(\frac{V_m + 40}{5}\right)}$ |
| $\alpha_{l,K,Adist}$ | $\exp\left(\frac{1.0e^{-3} \times 3 \times (V_m + 56) \times 9.648e^4}{8.315 \times (273.16 + C^\circ)}\right)$ |
| $Q_{10,K,Adist}$ | $5^{\frac{C^\circ - 24}{10}}$ |
| $a_{n,K,Adist}$ | 0.1 |
| $g_{HCN}$ | 0.0006 $S/cm^2$ |
| $E_{HCN}$ | -30 $mV$ |
| $\alpha_{l,HCN}$ | $\exp(0.0378 \times 4 \times (V_m + 90))$ if $dist_{soma} > 100\mu m$<br>$\exp(0.0378 \times 4 \times (V_m + 82))$ otherwise |
| $\beta_{l,HCN}$ | $\exp(0.0378 \times 0.88 \times (V_m + 75))$ |
| $\gamma_{l,HCN}$ | $\exp(0.0378 \times 2.2 \times (V_m + 75))$ |
| $Q_{10,HCN}$ | $4.5^{\frac{C^\circ - 33}{10}}$ |

#### Basket cells

Each basket cell has 89 compartments organized in somatic and dendritic groups. All compartments include a sodium current (*I*_*Na*_), a fast delayed rectifier potassium current (*I*_*K,fdr*_), an A-type potassium current (*I*_*K,A*_), a voltage-dependent calcium-activated potassium current (*I*_*K,vCa*_), a non-voltage-dependent calcium-activated potassium current (*I*_*K,Ca*_), a L-type calcium current (*I*_*Ca,L*_), a N-type calcium current (*I*_*Ca,N*_).

*I*_*Na*_ current is described by:

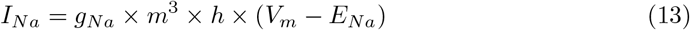

where *g*_*Na*_ is the maximum conductance, *E*_*Na*_ is the reversal potential and *m*, and *h* are gating variables defined by the equations:

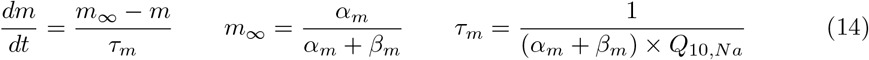

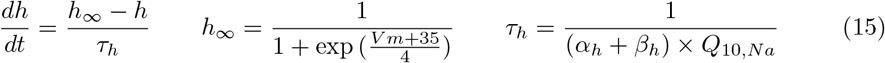

The potassium currents are described by:

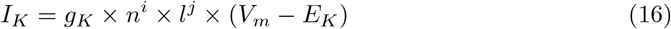

where *g*_*K*_ is the maximum conductance, *E*_*K*_ is the reversal potential, *n* and *l* are the gating variables defined by the equations:

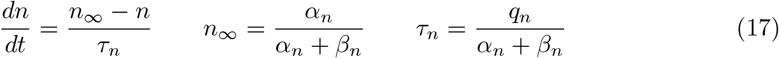

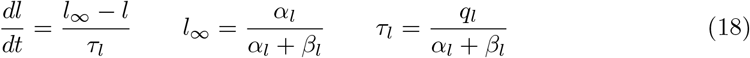

The calcium current *I*_*Ca,L*_ is described by:

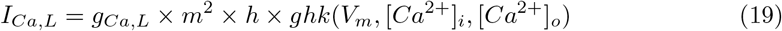

where *g*_*Ca,L*_ is the maximum conductance, *E*_*Ca,L*_ is the reversal potential, *m* and *h* are the gating variables defined by:

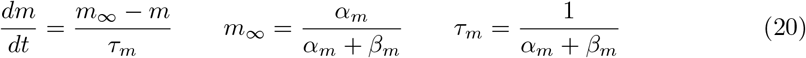

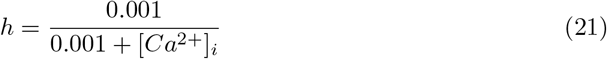

and *ghk* gives the driving force through the channel using the Goldman-Hodgkin-Katz equation:

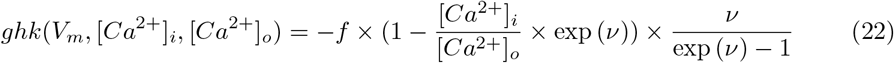

The calcium current *I*_*Ca;N*_ is described by:

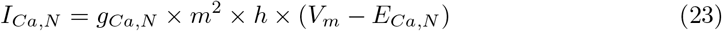

where *g*_*Ca,N*_ is the maximum conductance, *E*_*Ca,N*_ is the reversal potential, *m* and *h* are the gating variables defined by the equations:

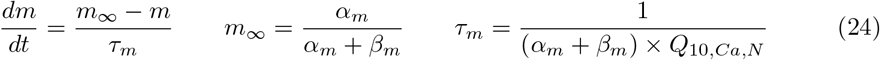

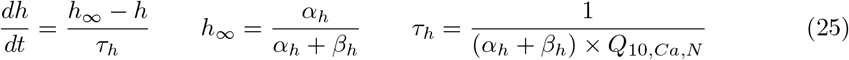

The full expressions for the parameters described above can be found in Table 2.

**Table 2.**
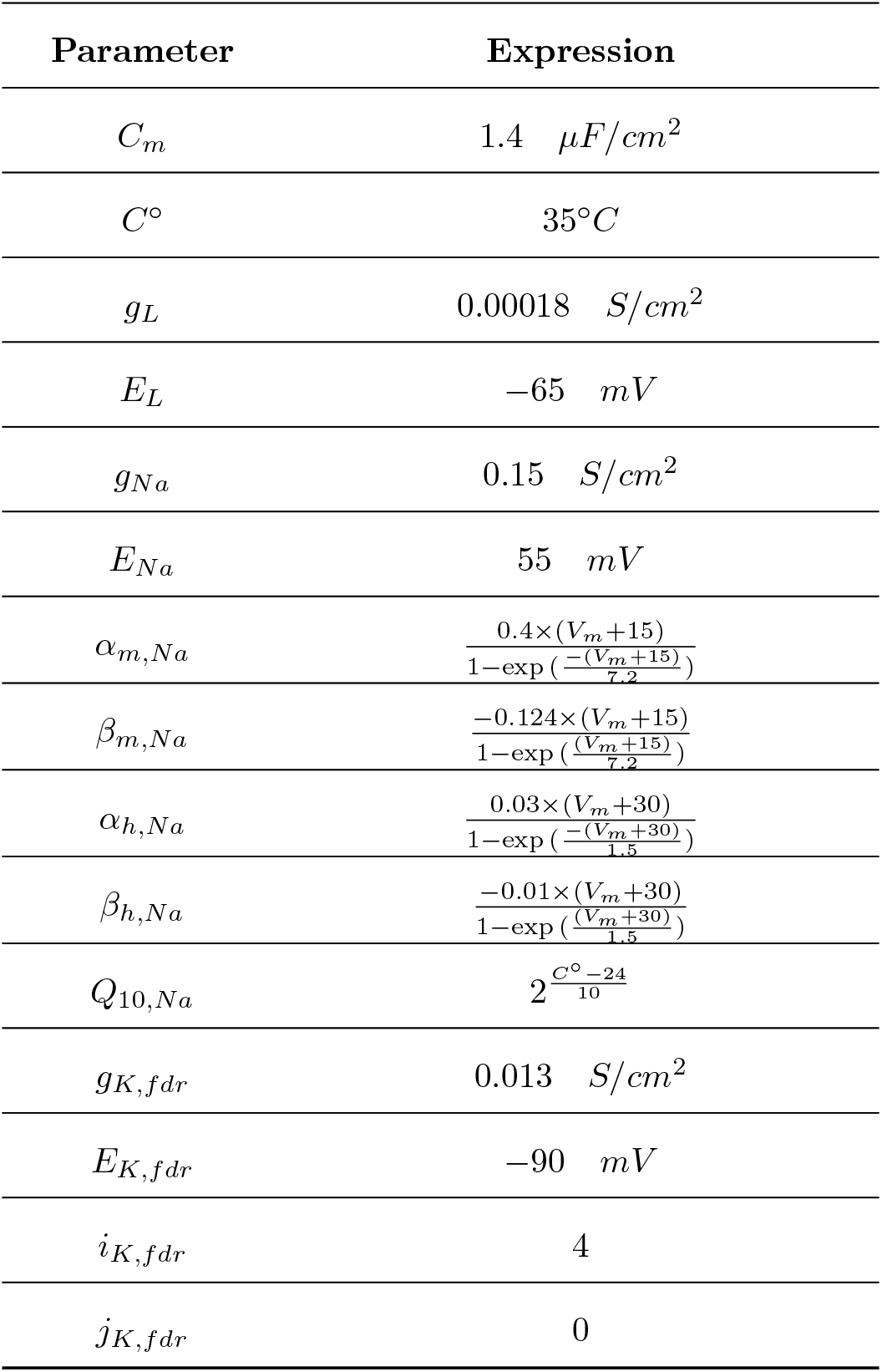
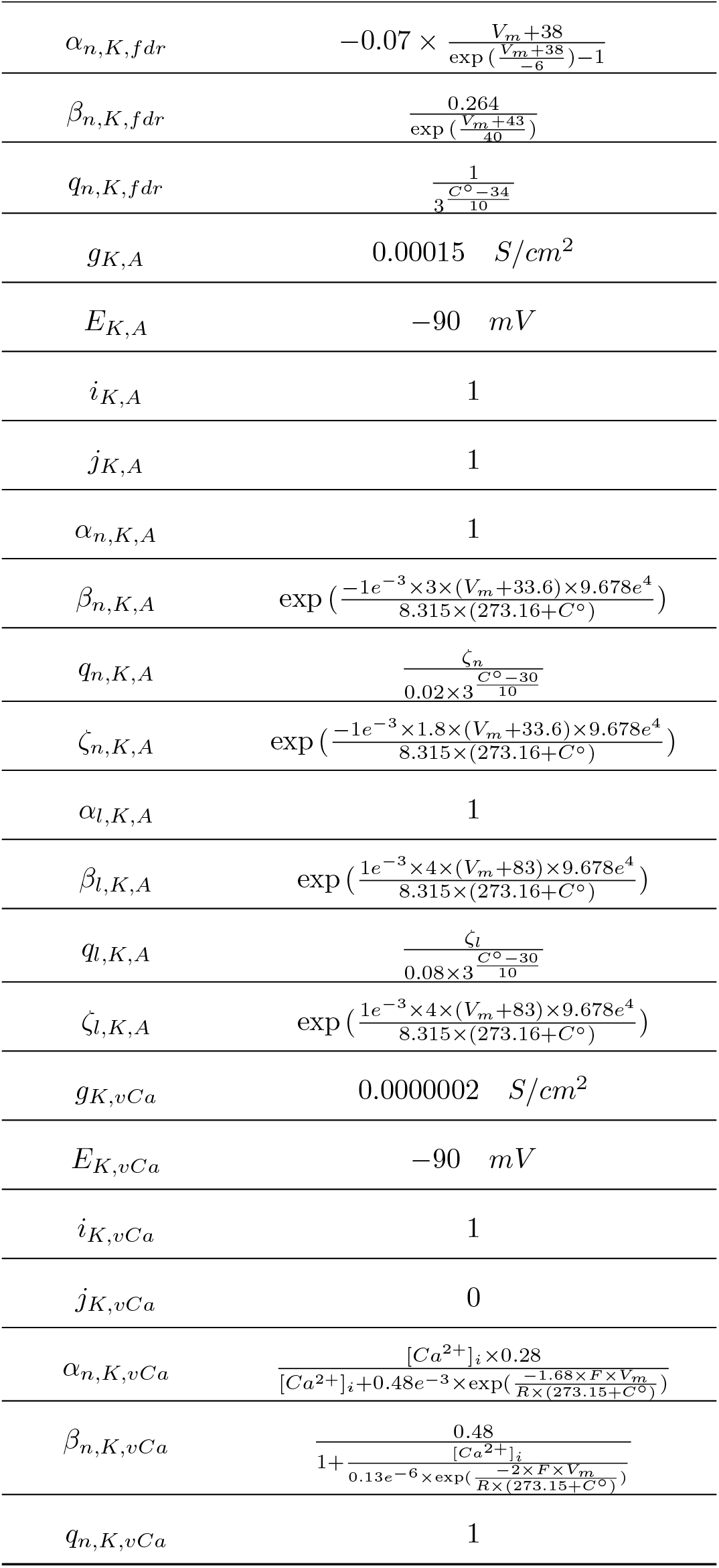
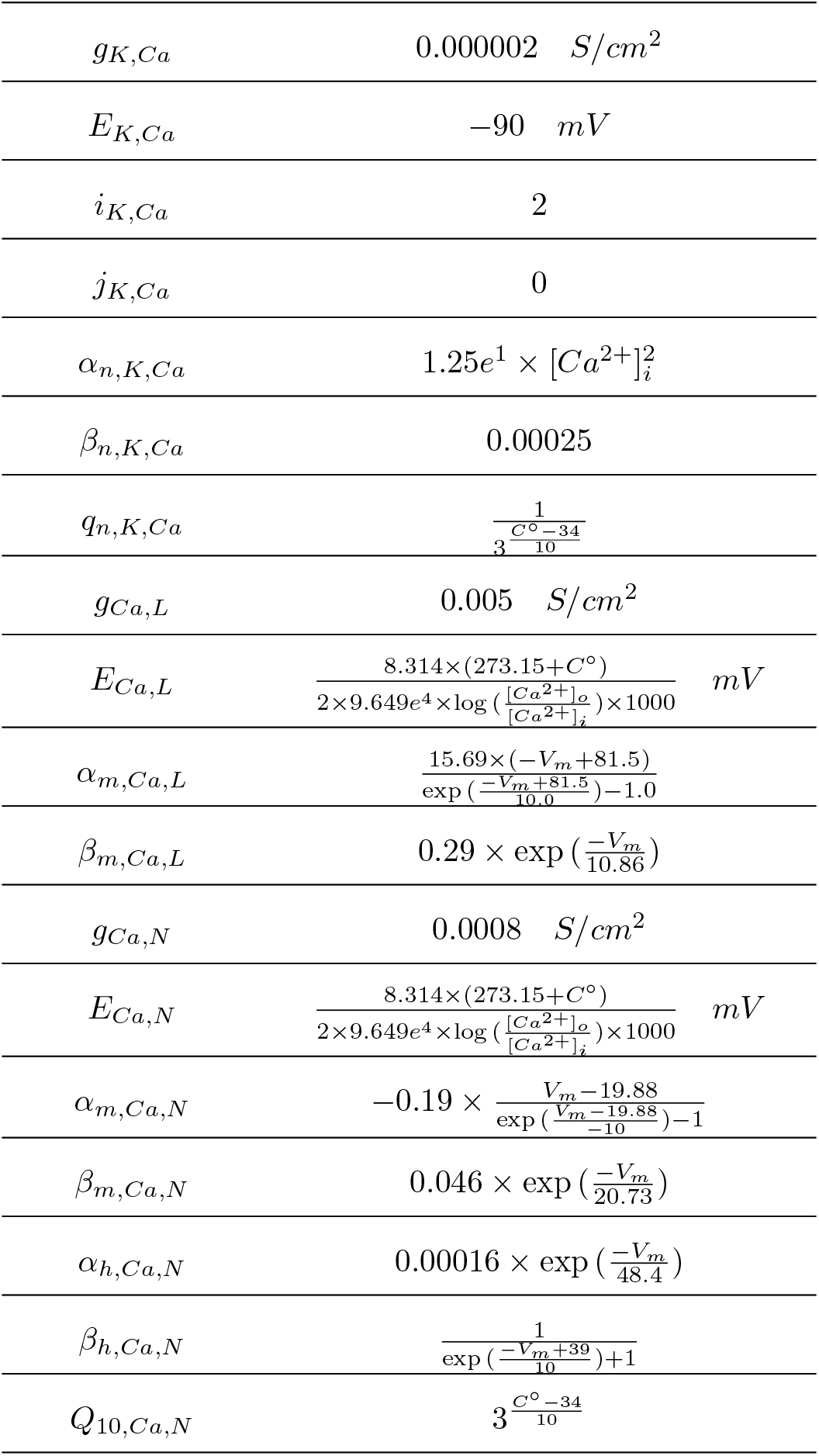
Parameter expressions for basket neurons.

#### OLM cells

OLM cells are represented with 34 compartments organized into somatic, dendritic and axonal groups. All compartments contain a sodium current (*I*_*Na*_) and a fast delayed rectifier potassium current (*I*_*K,fdr*_). The somatic and dendritic compartments also include an A-type potassium current (*I*_*K,A*_). In addition, the somatic compartments contain a hyperpolarization-activated cyclic nucleotide-gated current (*I*_*HCN*_).

The sodium current *I*_*Na*_ is described by:

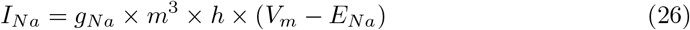

where *g*_*Na*_ is the maximum conductance, *E*_*Na*_ is the reversal potential, *m* and *h* are the gating variables defined by the equations:

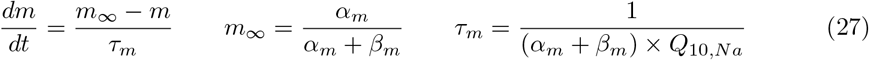

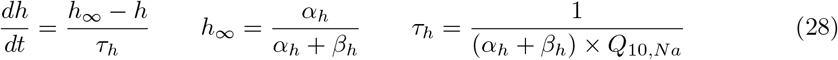

*I*_*K,fdr*_ current is described by:

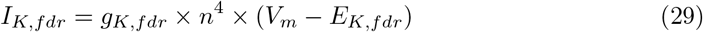

where *g*_*K,fdr*_ is the maximum conductance, *E*_*K,fdr*_ is the reversal potential and *n* is the gating variable defined by the equations:

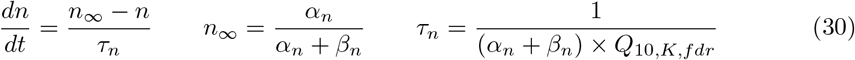

The potassium current *I*_*K,A*_ is described by:

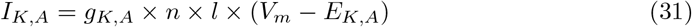

where *g*_*K,A*_ is the maximum conductance, *E*_*K,A*_ is the reversal potential, *n* and *l* are the gating variables defined by the equations:

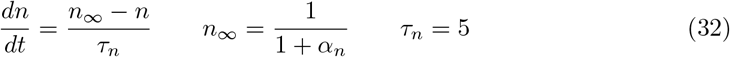

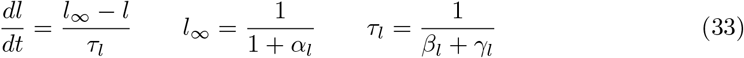

*I*_*HCN*_ current is described by:

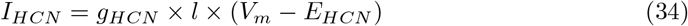

where *g*_*HCN*_ is the maximum conductance, *E*_*HCN*_ is the reversal potential and *l* is the gating variable defined by:

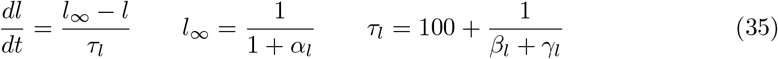

The full expressions for the parameters described above can be found in Table 3.

**Table 3.**
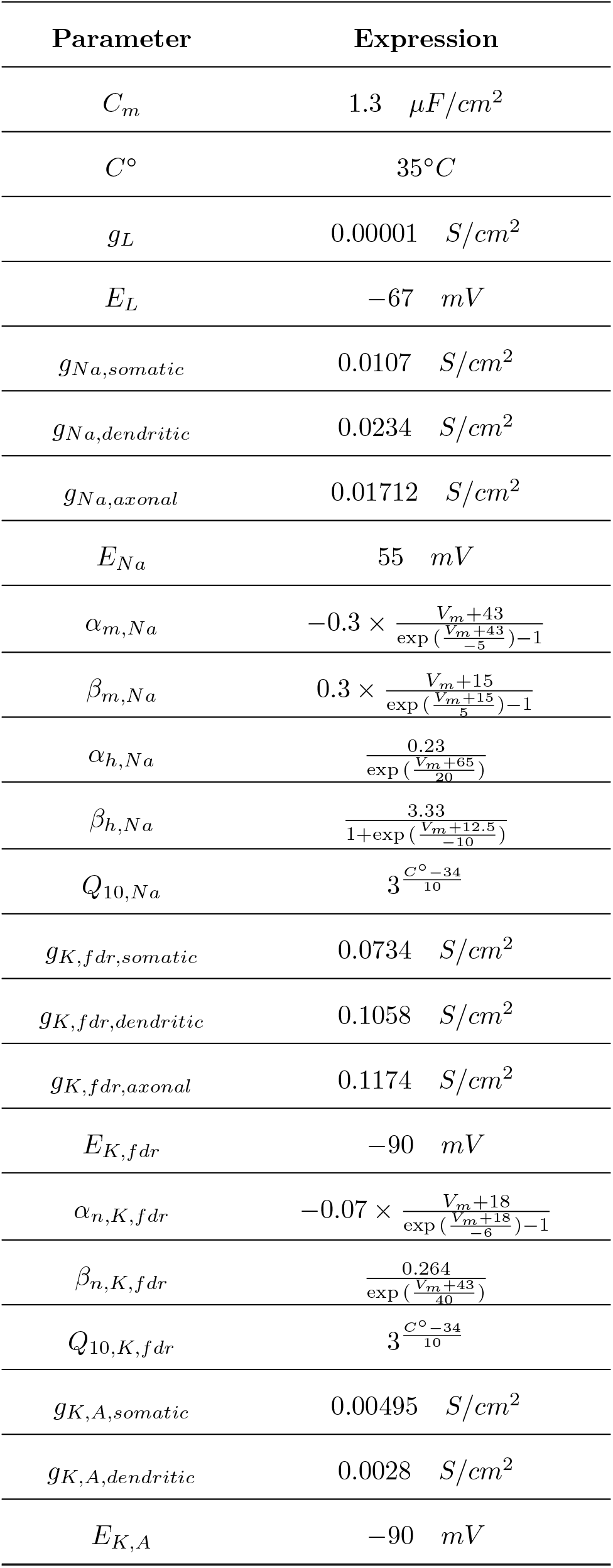
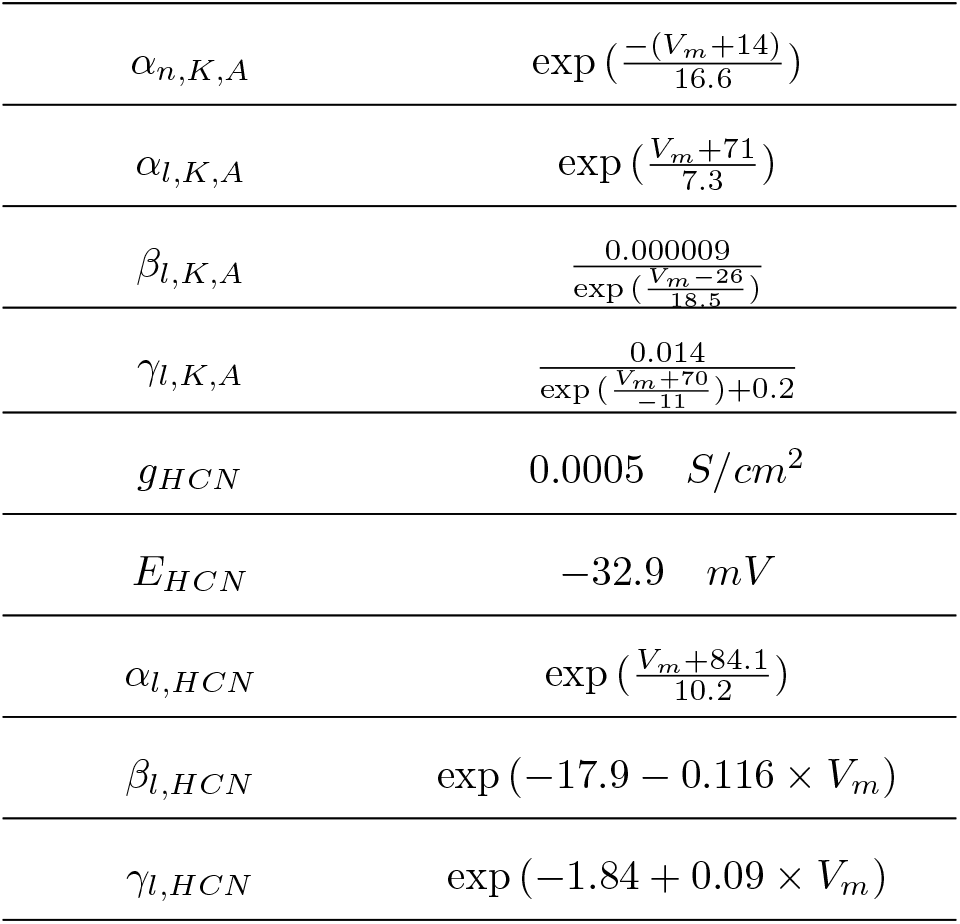
Parameter expressions for OLM neurons.

#### Schaffer collaterals

The number of compartments in each collateral depends on its length, ranging from a minimum of 45 to a maximum of 760 compartments in our model. Each internodal part of the collateral is subdivided into specialized compartments, as in [38]: one 1-µm-long node of Ranvier, two myelin sheath attachment segments (MYSA) of 3 µm each, two fluted region segments (FLUT) of 10 µm each, and six stereotyped internode segments (STIN) with a length of 15 µm each. This results in a total internode length of 117 µm. The modeled fibers were 2 µm in diameter. Each node of Ranvier compartment includes fast sodium (*I*_*Na,f*_), persistent sodium (*I*_*Na,p*_), slow potassium (*I*_*Ks*_) and leak (*I*_*L*_) currents.

The sodium currents are described by:

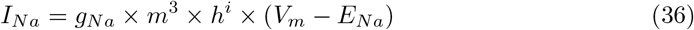

where *g*_*Na*_ is the maximum conductance, *E*_*Na*_ is the reversal potential, *m* and *h* are the gating variables defined by:

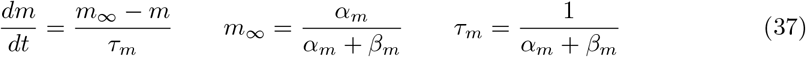

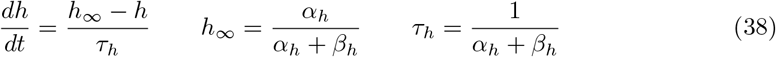

The potassium current is described by:

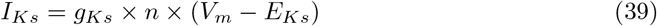

where *g*_*Ks*_ is the maximum conductance, *E*_*Ks*_ is the reversal potential and *n* is the gating variable defined by:

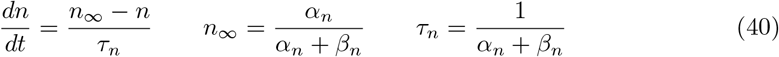

The full expressions for the parameters described above can be found in Table 4.

**Table 4.** Parameter expressions for Schaffer collaterals’ node of Ranvier.

| Parameter | Expression |
| --- | --- |
| $C_m$ | $2 \mu F/cm^2$ |
| $C^\circ$ | $35^\circ C$ |
| $g_L$ | $0.007 S/cm^2$ |
| $E_L$ | $-90 mV$ |
| $g_{Na,f}$ | $3 S/cm^2$ |
| $E_{Na,f}$ | $50 mV$ |
| $i_{Na,f}$ | $1$ |
| $\alpha_{m,Na,f}$ | $Q_{10,m} \times \frac{1.86 \times (V_m + 21.4)}{1 - \exp\left(\frac{-(V_m + 21.4)}{10.3}\right)}$ |
| $\beta_{m,Na,f}$ | $Q_{10,m} \times \frac{-0.086 \times (V_m + 25.7)}{1 - \exp\left(\frac{V_m + 25.7}{9.16}\right)}$ |
| $\alpha_{h,Na,f}$ | $Q_{10,h} \times \frac{-0.062 \times (V_m + 114)}{1 - \exp\left(\frac{V_m + 112}{11}\right)}$ |
| $\beta_{h,Na,f}$ | $\frac{Q_{10,h} \times 2.3}{1 + \exp\left(\frac{-(V_m + 31.8)}{13.4}\right)}$ |
| $g_{Na,p}$ | $0.01 S/cm^2$ |
| $E_{Na,p}$ | $50 mV$ |
| $i_{Na,p}$ | $0$ |
| $\alpha_{m,Na,p}$ | $Q_{10,m} \times \frac{0.3 \times (V_m + 27)}{1 - \exp\left(\frac{-(V_m + 27)}{10.2}\right)}$ |
| $\beta_{m,Na,p}$ | $Q_{10,m} \times \frac{-0.00025 \times (V_m + 34)}{1 - \exp\left(\frac{V_m + 34}{10}\right)}$ |
| $Q_{10,m}$ | $2.2^{\frac{C^\circ - 20}{10}}$ |
| $Q_{10,h}$ | $2.9^{\frac{C^\circ - 20}{10}}$ |
| $g_{Ks}$ | $0.08 S/cm^2$ |
| $E_{Ks}$ | $-90 mV$ |
| $\alpha_n$ | $\frac{Q_{10,n} \times 0.3}{1 + \exp\left(\frac{V_m + 53}{-5}\right)}$ |
| $\beta_n$ | $\frac{Q_{10,n} \times 0.03}{1 + \exp\left(\frac{V_m + 90}{-1}\right)}$ |
| $Q_{10,n}$ | $3^{\frac{C^\circ - 36}{10}}$ |

## Model architecture

### Number of neurons

The number of principal neurons present in the CA1 area in the human brain is on the order of 14 million [64]. However, to mitigate the computational cost, we reduced the number of neurons used in our model, while keeping the ratios between the different cell types. For instance, interneurons are known to represent about 8-10% of the neurons in hippocampal regions. Furthermore, PV+ basket cells constitute around 14.4% of the interneurons in hippocampal regions, while OLM cells make up for about 4.3% of them [65], which can be approximated to a ratio of 3:1 between basket cells and OLM cells. A population of 100 pyramidal cells was therefore used to allow the emergence of stable collective dynamics while maintaining a computationally tractable model. We modeled 9 basket cells and 3 OLM cells to keep the ratio between the cell types.

Furthermore, a minimum of 3 OLM cells was required to achieve synaptic coverage across the entire CA1 area (see the section below on neuronal connections).

We also wanted to represent explicit projections of Schaffer collaterals, which are axonal projections from CA3 pyramidal cells to the CA1 area. There are about 2.8 million pyramidal cells in the CA3 area of the human brain [64]. We approximated the ratio between CA1 and CA3 pyramidal cells to 4:1, thus modeling 26 Schaffer collaterals.

The number of neurons represented for each cell type is summarized in Table 5.

**Table 5.** Network parameters.

|  | Pyramidal cell | Basket cell | OLM cell | Schaffer collaterals |
| --- | --- | --- | --- | --- |
| Number of neurons | 100 | 9 | 3 | 26 |
| Synaptic distance (μm) | 500 | 470 | 1057.695 | 309.165 |

### Geometry of the model

The hippocampal formation has a curved shape, with excitatory neurons projecting along its proximo-distal axis. The entorhinal cortex lies at the most distal end of this axis and the dentate gyrus forms the proximal end (see Fig. 1C). To accurately represent the trajectory of Schaffer collaterals from CA3 to CA1, as well as the overall organization of CA1, we used an intrinsic coordinate system (*t, u*) in our model, where the *t*-axis corresponds to the curvilinear axis and the *u*-axis represents the radial, superficial-to-deep axis (from stratum oriens to stratum lacunosum moleculare as seen in Fig. 1D).

To achieve this, we simplified the topology of the hippocampal formation slice using a generalized S-shape, extracted from the Hippocampus book [66]. This shape has the advantage of connecting every region one after the other, allowing the *t*-axis to remain continuous. As a result, spatial manipulations and distance computations become more straightforward. For example, the curved structure can be easily transformed into a flattened representation for better interpretability. We manually defined (using Inkscape) two cubic Bézier curves joined end to end, following the generalized S-shape. Each Bézier curve is defined by the following equation:

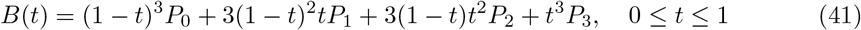

where *P*_0_, *P*_1_, *P*_2_, *P*_3_ are control points which provide directional information on the curves. *t* is the curves’ parameter, normalized between 0 and 1, indicating the position along the curve, with 0 being the starting point of the curve and 1 being the ending point. We stacked the two curves together to use the same evolving *t* ∈ [0, 1], resulting in 7 control points (*P*_0_, *P*_1_, *P*_2_, *P*_3_, *P*_4_, *P*_5_ and *P*_6_) and the following piecewise equation:

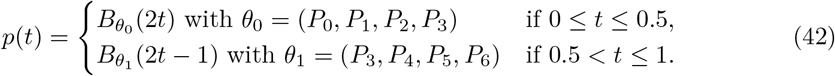

Our S-curve was first constructed by sampling *n*_*points*_ = 100 points at uniformly spaced intervals (*t* = 0, 0.01, 0.02, …, 1.0) along the piecewise Bézier curve defined in equation (42). However, these points were not evenly spaced along the curve as Bézier parametrization does not have a constant acceleration. An additional mapping operation was applied to redistribute the points so that the spacing between their corresponding *t* values reflect uniform distances along the curve length. In this mapping, each parameter *t* was recalculated so that a given value, *t* = 0.82 for example, represents the point located at 82% of the total curve length. In order to do that, we computed the cumulative arc length from the starting point of the curve to each parameter value *t* and stored each resulting value in a list *arc*_*lengths*. We also calculated the total arc length of the curve *L*_*total*_. Then, for each parameter *t*, we determined the corresponding desired arc length *a*(*t*):

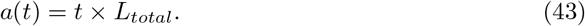

Next, we searched for the index *i* that has the closest value between *arc*_*lengths*_*i*_ and *a*(*t*). If *arc*_*lengths*_*i*_ was bigger than the value of *a*(*t*), we decreased the index *i* by 1 (*i* = *i* − 1). We finally determined the mapping *t* of each parameter *t* as follows :

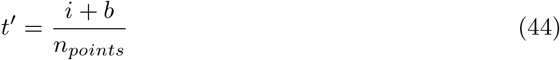

with :

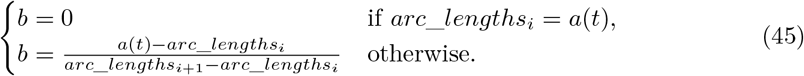

The resulting piecewise equation describing the Bézier curve is:

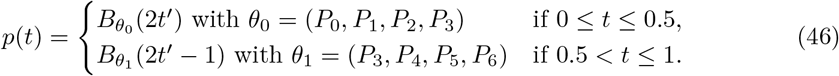

The width of our hippocampal slice was fixed to *l*_*shape*_ = 1.5 mm, consistent with dimensions from human MRI data as reproduced in the work of Aussel and colleagues [30]. The *u*-axis represents the normalized width of the slice.

From the intrinsic coordinates, we can get the unfolded coordinates (*t*_*unfold*_, *u*_*unfold*_) by multiplying the *t*-coordinate by the total curvilinear length (*L*_*total*_) and the *u*-coordinate by the total width of the slice (*l*_*shape*_). We used the unfolded coordinates for neuron connection.

We can also retrieve the corresponding Cartesian coordinates (*x, y*) in the following way. The coordinates (*x*_*p*_, *y*_*p*_) = *p*(*t*) correspond to the intrinsic coordinates (*t*, 0.5), as the Bézier curve intersects the *u*-axis perpendicularly at its midpoint. To obtain the coordinates (*x, y*) corresponding to (*t, u*), we firsr determine the slope *m* of the tangent vector to the Bézier curve at the point *p*(*t*) and then compute:

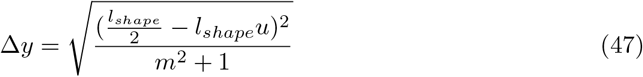

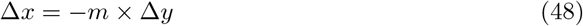

Finally, the Cartesian coordinates are obtained by:

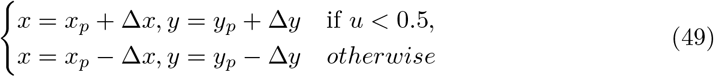

#### Neuron placement

Neuron placement was assigned using the intrinsic coordinate system (*t, u*) described in the previous subsection. The *u*-coordinate of each CA1 neuron’s soma was constrained by the anatomical boundaries of its respective layer: OLM cells were placed within the stratum oriens, while pyramidal and basket cells soma were restricted to the stratum pyramidale (see Fig. 1D). Due to the relatively small number of inhibitory neurons, their *t*-coordinates were chosen deterministically to ensure even spacing along the curvilinear axis. This was achieved by dividing the transverse length of the CA1 area by the number of neurons plus one, thereby avoiding placement directly on the region’s boundaries. On the other hand, the *t*-coordinates of pyramidal neuron somata were assigned randomly using a uniform distribution along the curvilinear axis of the CA1 region. To ensure functional inhibitory connectivity, an additional constraint was applied: each pyramidal neuron had to lie within 470 µm of a basket cell, guaranteeing that it received inhibitory input from at least one basket cell (see next section).

To place the Schaffer collaterals, we first determined the coordinates of the first node for each collateral. As with the CA1 pyramidal neurons, the *u*-coordinates were sampled from a uniform distribution within the stratum pyramidale. The *t*-coordinates were randomly assigned using a uniform distribution along the curvilinear axis of the CA3 region. Next, we assigned a length to each collateral (*L*_*SCA*_). To do this, we divided the CA3 region into two equal parts: a proximal CA3 half, which borders the dentate gyrus, and a distal half, which borders the CA1 region. Similarly, the CA1 region was divided into two halves: a proximal CA1 half, adjacent to the CA3 area, and a distal CA1 half, which borders the subiculum. Schaffer collaterals arising from pyramidal cells in the proximal CA3 region project to distal portions of the CA1 area, near to the subiculum. Conversely, collaterals originating from distal CA3 neurons terminate in proximal CA1, adjacent to the CA3 area [66]. Therefore, the length of each collateral can be calculated as follows:

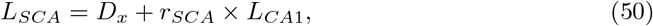

where *D*_*x*_ is the intrinsic distance of the collateral’s first node to the border between CA3 and CA1 regions; *r*_*SCA*_ is a random number between 0.5 and 1 if the first node is located in the proximal CA3, and between 0 and 0.5 otherwise; *L*_*CA*1_ is the curvilinear length of the CA1 area. Given the coordinates of the first node and the total length of the collateral, we generated the trajectory of each collateral using Bézier curves, as described in the previous section. Unlike the modeled CA1 neurons, which had a fixed topology and a fixed number of compartments, the number of compartments of each Schaffer collateral depends on its length. In our model, this number ranges from 45 to 760 compartments per collateral. However, the length of each compartment was kept the same as in [38].

#### Neuronal connectivity

Pyramidal neurons form synapses on the soma of both basket and OLM cells. There are few connections from pyramidal to pyramidal neurons in the CA1 area. On average, one pyramidal neuron connects to 1% of pyramidal neurons [67]. We compared simulations in which pyramidal neurons synapsed onto exactly one other pyramidal neuron with simulations that excluded pyramidal-to-pyramidal connections, and found no significant differences in network behaviour. Therefore, no pyramidal-to-pyramidal connection was included in the final model.

As for the synaptic connections included in the model, basket cells synapse on the soma of pyramidal neurons and other basket cells. OLM cells inhibit the apical tuft of the pyramidal neurons, which corresponds to the distal apical dendrites located in the stratum lacunosum moleculare. In our model, this region is represented by a list of 6 dendritic sections. For each OLM-to-Pyramidal connection, one section was randomly selected from this list as the synapse target. Pyramidal cells also receive excitatory input from the Schaffer collaterals onto their apical trunk, modeled as 3 dendritic sections located in the stratum radiatum (see Fig. 1B). One section was randomly selected as the target for each Schaffer collateral input.

The connectivity in our model is distance-based, meaning that neurons form synapses with other neurons in close proximity. The Hippocampome.org[68] database provides the mean axonal length, which we will refer to as “synaptic distance”, for each cell type. To account for anatomical scale differences between mouse and human brains, we scaled the database values by multiplying them by a factor of 1.5. Next, we created histograms to visualize the distances between each cell type and their potential targets in our model, and overlaid the synaptic distances on top of these histograms in order to validate this approach. This revealed that no basket cell was within synaptic distance of another basket cell, likely due to the small number of basket cells in the model. To address this, we increased their synaptic distance as to get basket-to-basket connections. Similarly, we increased the synaptic distance for pyramidal neurons to enhance the pyramidal-to-OLM connections, so as to get all the OLM cells receive connections from the network. Finally, each cell was connected to any other cell that was both within its synaptic distance and belonged to one of its target cell types.

For the Schaffer collaterals, which synapse onto pyramidal cells in the stratum radiatum (see Fig. 1B), we took the mean dendritic length of pyramidal cells in that layer from the database and scaled it by a factor of 1.5 to use as the synaptic distance. Schaffer collaterals were connected to any pyramidal cell whose soma laid within that distance from their last node. The synaptic distance values for each cell type are summarized in Table 5.

### Synapses

We considered AMPA and GABA-A synapses modeled using the double exponential synapse mechanism (Exp2Syn) provided by the NEURON simulator [69], which is characterized by the following equations :

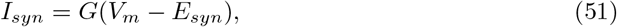

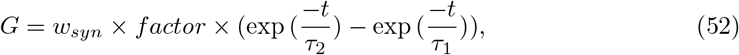

where *I*_*syn*_ is the synaptic current, *E*_*syn*_ represents the synaptic reversal potential, *w*_*syn*_ is the synaptic weight, *τ*_1_ and *τ*_2_ are time constants corresponding to the rise and decay phases, respectively. The synaptic parameters can be found in Table 6.

**Table 6.**
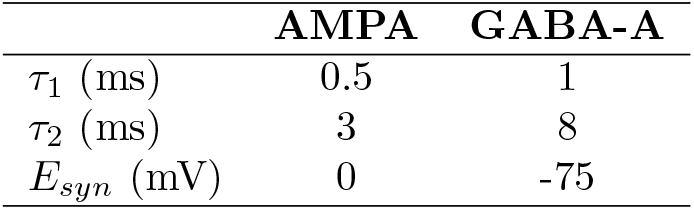
Synaptic parameters.

To determine the synaptic connection weights between cell types, we first considered the network in the absence of inputs from Schaffer collaterals, as illustrated in Fig. 2A. Our first approach consisted in applying the method of Romaro and colleagues [52] to the work of Vardalakis and colleagues [32]. Since the model used by Vardalakis and colleagues [32] did not include OLM cells, we assigned the same synaptic weights to basket cells and OLM cells. However, we were not able to get gamma activity. This is probably due to the small size of our model. In addition, whereas the model developed by Vardalakis and colleagues relied on single-compartment neurons, our study employed multicompartment neuron models, which may require further parameter tuning to achieve comparable dynamics.

Therefore, we adopted another strategy to tune the connection weights. First, we set all inhibitory and excitatory synaptic weights to zero. Then, we injected a 6 Hz oscillatory input with an amplitude of 6 nA into the apical dendrites of each pyramidal neuron to mimic excitatory input from the entorhinal cortex. We incrementally increased the strength of excitatory connections until all three populations exhibited sufficient activities. The resulting weights were used as baseline values for excitatory synaptic weights (denoted *w*_*EBC*_ and *w*_*EOLM*_ in Fig. 2A). For the inhibitory connections, we set the baseline weights (*w*_*IE*_ in Fig. 2A) to a value scaled from the inhibitory weights used in [32]. The initial weights are listed in Table 7.

**Table 7.** Initial synaptic weights (µS)

| From \ To | Pyramidal | Basket | OLM | Schaffer collaterals |
| --- | --- | --- | --- | --- |
| **Pyramidal** | - | 0.0045 | 0.0009 | - |
| **Basket** | 0.36 | 0.0018 | - | - |
| **OLM** | 0.36 | - | - | - |
| **Schaffer collaterals** | 0.009 | - | - | - |

Once all the baseline synaptic weights were established, we performed a grid search to optimize the balance between excitation and inhibition in order to get gamma-band activity in all cell populations (Fig.2). We injected an intracellular current ramp, with an amplitude ranging from 0 to 1 nA, into the soma of all pyramidal cells. We conducted several 5-second simulations during which we varied two scaling factors independently:

- *k*_*E*_: scaling factor for excitatory synaptic weights from pyramidal cells to basket and OLM cells (ranging from 0.1 to 2)
- *k*_*I*_ : scaling factor for inhibitory weights from both interneuron populations to pyramidal cells (ranging from 0.1 to 1.0)

We measured the mean oscillatory frequency of each cell population during the last second of simulation. For each cell type, we generated a heatmap showing their oscillatory frequency as a function of *k*_*E*_ and *k*_*I*_ (see Fig. 2B). We also computed a heatmap showing the ratio between pyramidal and basket cells oscillatory frequencies over the last second of simulation (see Fig. 2C). Our goal was to achieve a minimum oscillatory frequency of 30 Hz in all populations and to maintain a ratio of at least 2 between the basket and pyramidal populations firing rates as pyramidal cells discharge at a lower rate than basket cells [48]. Based on these criteria, we selected *k*_*E*_ = 2 and *k*_*I*_ = 0.1 (which are indicated by white dashed lines in the panels B and C of Fig. 2). The resulting network activity shown in Fig. 2D, confirmed that all populations reached oscillatory frequencies above 30 Hz towards the last seconds of simulation. The fixed synaptic weights used for each connection type are summarized in Table 8.

**Table 8.** Fixed synaptic weights (µS)

| From \ To | Pyramidal | Basket | OLM | Schaffer collaterals |
| --- | --- | --- | --- | --- |
| **Pyramidal** | - | 0.009 | 0.0018 | - |
| **Basket** | 0.036 | 0.0018 | - | - |
| **OLM** | 0.036 | - | - | - |
| **Schaffer collaterals** | 0.0261 | - | - | - |

Once the synaptic weights between cell types were fixed, we incorporated the Schaffer collaterals into the network, as depicted in Fig. 3A. The first node of each Schaffer collateral (i.e. the endpoint located in CA3) received a 6 Hz oscillatory input. An offset randomly sampled between 0 and 50 ms was added to the onset of theta inputs for additional noise. The Schaffer collaterals synapse onto the trunk of pyramidal cells. We observed that both the amplitude of the oscillatory input to Schaffer collaterals and synaptic weight of Schaffer collaterals to pyramidal cells influenced the network dynamic in CA1. To optimize the synaptic weight from the CA3 Schaffer collaterals to the CA1 pyramidal population (denoted *w*_*SCA*_ in Fig. 3A), we initialized *w*_*SCA*_ to the same tuned value as *w*_*EBC*_ = 0.009 (see Table 8). We performed multiple 5-second simulations in which we independently varied:

- *k*_*SCA*_: scaling factor for Schaffer collateral-to-pyramidal cell synaptic weight (ranging from 2 to 4)
- *k*_*amp*_: amplitude of the theta input (ranging from 0.02 to 0.2 nA)

The mean oscillatory frequencies of all populations within the phases of the ongoing theta were measured. We generated heatmaps showing the oscillatory frequencies of pyramidal and basket cells as a function of *k*_*amp*_ and *k*_*SCA*_ (see Fig. 3B). The aim was to maximize the firing activity of both populations. As a result, we selected *k*_*amp*_ = 0.1 nA and *k*_*SCA*_ = 2.9, as indicated by orange dashed lines in Fig. 3B. The resulting network activity shown in Fig. 3C exhibited mean oscillatory activity above 30 Hz for OLM, basket and pyramidal cells, which were synchronized to the ongoing theta cycle.

### Extracellular stimulation

In addition to the intracellular current applied to Schaffer collaterals (which represents the external inputs to the CA1 area) presented above, we applied extracellular stimulation to model the effects of hippocampal DBS. Extracellular stimulation was modeled under the assumption of a fully resistive and isotropic infinite extracellular medium. Furthermore, the currents generated by a neuron’s electrical activity had no effect on the stimulating field. The value of the extracellular potential measured at each compartment (*V*_*e,n*_) of a neuron is determined by:

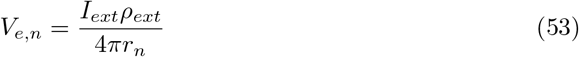

for a monopolar stimulation. *I*_*ext*_ represents the amplitude of the injected extracellular current. *ρ*_*ext*_ is the resistivity of the extracellular medium and *r*_*n*_ is the distance of the compartment to the center of the electrode.

We also considered bipolar stimulations, represented by the following equation:

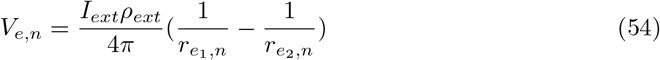

where 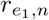 and 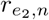 are the distances of the compartment to the middle of the first and the second electrode respectively.

When applying extracellular stimulation to the full CA1 network, we considered a minimum distance of 400 µm between neuronal compartments and electrode center, with a 1.5 mm separation between the two electrodes for bipolar stimulation. This configuration was chosen to match the physical dimensions of stereoencephalography (sEEG) electrodes, which have a 800 µm diameter and a 1.5 mm inter-electrode distance. We also neglected the presence of neurons located between the electrode and the target compartment, as well as the extracellular electric fields generated by the activity of neighboring neurons, when computing the extracellular potential.

#### Stimulation parameters

Various sets of stimulation parameters were employed. To generate Fig. 4, we applied a 1-ms single rectangular pulse after an offset of 100 ms. The stimulation amplitude varied from -10 mA to 10 mA.

For the biphasic stimulations shown in Figs. 5-8, we used a pulse train consisting of anodic and cathodic rectangular pulses. The pulse width was set to 300 µs with an interphase of 100 µs. Amplitudes ranged from 0.5 to 6 mA for Figs. 5 and 6, and were fixed at 3.5 mA for Figs. 7 and 8, using stimulation frequencies of either 50 Hz of 130 Hz. The stimulation was initiated after a 2 s delay and lasted for 2 s.

To produce Fig. 7C, theta-burst stimulation was applied to the network. The protocol consisted of a 5 Hz burst frequency, where each burst comprised a series of anodic and cathodic pulses at 100 Hz. The pulse width was set at 200 µs with an interphase of 100 µs. The amplitude was set to 3.5 mA. Stimulation was initiated at 2 s of simulation time and lasted for 2 s.

#### Electrodes configurations

We tested several configuration of electrodes. To determine the electrode coordinates, we used the intrinsic coordinate system.

For monopolar electrodes, the *t*-coordinate was set to the middle of CA1 curvilinear length. For the electrode placed on the outer curvature, the *u*-coordinate was set to -0.5. For the electrode placed on the inner curvature, *u* = 1.5.

For the bipolar electrodes oriented perpendicularly to the Schaffer collaterals, *t*_1_ and *t*_2_ were set to the middle of CA1 curvilinear length. For the electrodes placed on the outer curvature, *u*_1_ = -0.5 and *u*_2_ = -1.5. For the electrodes placed on the inner curvature, *u*_1_ = 1.5 and *u*_2_ = 2.5.

For the electrodes oriented parallel to the Schaffer collaterals, the positions were set to 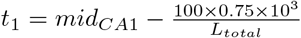 and 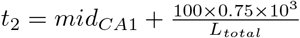, placing each electrode 0.75 mm away from the CA1 midpoint along the curvilinear axis, resulting in an inter-electrode spacing of 1.5 mm. For the electrodes placed on the outer curvature, *u*_1_ = *u*_2_ = -0.5. For the electrodes placed on the inner curvature, *u*_1_ = *u*_2_ = 1.5.

### Data analysis

For each neuron and Schaffer collateral, we monitored its spike timing and its membrane potential. The data were recorded in the soma for pyramidal, basket and OLM cells, and in the last node for the Schaffer collaterals (i.e. the endpoint located in CA1). The threshold for spike detection was set to 0 mV.

#### Firing rates

To compute the firing rate of each population, spike times were segmented into 5-ms windows with 90% overlap. The spike count within each window was normalized by the window duration and the population size (Fig. 2B).

#### Spectrograms

Spectrograms of the binned firing rates were computed using the short-time Fourier transform with a sliding Hann window of 150 ms width and 90% overlap (Fig. 2D and Fig. 3C).

#### Dominant oscillatory frequency

From the spectrograms, we computed the dominant oscillatory frequencies *f*_*dominant*_ for each population. They were calculated as the mean power-weighted frequency within each theta phase:

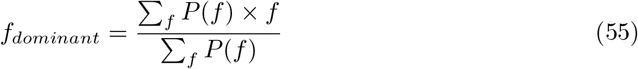

where *f* is the frequency in Hz and *P* (*f*) is the power at frequency *f* within a theta cycle. The frequency band was delimited between 0 and 150 Hz. To generate the heatmaps in Fig. 3B, we computed *f*_*dominant*_ of each theta cycle, starting at 2 s of simulation to the end of the simulation. Then, we calculated the mean of the oscillatory frequencies.

#### Change in spiking activity under extracellular stimulation

To determine the change in spiking activity in each population during extracellular stimulation (Fig. 5B, Fig. 6B, Fig. 7E and Fig. 8E), we divided the spike times into 166-ms windows with no overlap, starting at the onset of the oscillatory inputs. Each window therefore corresponded to one theta cycle. The spike counts outside the stimulation span (before 2 s and after 4 s of simulation) served as baseline activity. For each population, the spike count per neuron was measured during each window. We computed the mean spike count, as well as the standard deviation, for each neuron over all the windows for baseline activity. Following the same principle, we also computed the mean number of spikes for each neuron during the stimulation span (between 2 s and 4 s). Then, we considered that a neuron had an increased activity during the stimulation span if its mean spike count during stimulation was higher than its mean spike count plus standard deviation during baseline activity. Similarly, we considered that a neuron had a decreased activity during the stimulation span if its mean spike count during stimulation was smaller than its mean spike count minus standard deviation during baseline activity. We computed the number of neurons that had an increase or a decrease in activity for each neuronal population and we divided it by the population size.

### Simulation setup and code availability

The model was implemented using NEURON 8.2 with Python 3.10.9 [63]. Simulations were performed using a timestep of 0.025 ms. The total duration of simulations varied between 350 ms for single-cell stimulation amplitude threshold and 5 s for the other experiments (see Results). Simulations for weights and input optimization, as well as stimulation amplitude threshold were carried out using the Grid’5000 testbed, supported by a scientific interest group hosted by Inria and including CNRS, RENATER and several Universities as well as other organizations (see https://www.grid5000.fr). All other simulations were performed on a PC running on Windows 11. The average time for simulating 5 s of the full network locally was 15 min, and 11 min in the absence of Schaffer collaterals.

All model files and analysis scripts are available at the following Github repository under the GNU General Public License v3.0 license: https://github.com/Mandriantso/multicomp_hipp.

## Acknowledgments

This project was funded by the Inria PhD Program. Experiments presented in this article were carried out using the Grid’5000 testbed, supported by a scientific interest group hosted by Inria and including CNRS, RENATER and several Universities as well as other organizations (see https://www.grid5000.fr).

## References

1. Rolls ET. A computational theory of episodic memory formation in the hippocampus. Behavioural Brain Research. 2010;215(2):180–196. doi:10.1016/j.bbr.2010.03.027.

2. Danieli K, Guyon A, Bethus I. Episodic Memory formation: A review of complex Hippocampus input pathways. Progress in Neuro-Psychopharmacology and Biological Psychiatry. 2023;126:110757. doi:10.1016/j.pnpbp.2023.110757.

3. Colgin LL. Theta–gamma coupling in the entorhinal–hippocampal system. Current Opinion in Neurobiology. 2015;31:45–50. doi:10.1016/j.conb.2014.08.001.

4. Colgin LL. Rhythms of the hippocampal network. Nature Reviews Neuroscience. 2016;17(4):239–249. doi:10.1038/nrn.2016.21.

5. Axmacher N, Henseler MM, Jensen O, Weinreich I, Elger CE, Fell J. Cross-frequency coupling supports multi-item working memory in the human hippocampus. Proceedings of the National Academy of Sciences. 2010;107(7):3228–3233. doi:10.1073/pnas.0911531107.

6. Fell J, Axmacher N. The role of phase synchronization in memory processes. Nature Reviews Neuroscience. 2011;12(2):105–118. doi:10.1038/nrn2979.

7. Tort ABL, Komorowski RW, Manns JR, Kopell NJ, Eichenbaum H. Theta–gamma coupling increases during the learning of item–context associations. Proceedings of the National Academy of Sciences. 2009;106(49):20942–20947. doi:10.1073/pnas.0911331106.

8. Canolty RT, Knight RT. The functional role of cross-frequency coupling. Trends in Cognitive Sciences. 2010;14(11):506–515. doi:10.1016/j.tics.2010.09.001.

9. Lega B, Burke J, Jacobs J, Kahana MJ. Slow-Theta-to-Gamma Phase–Amplitude Coupling in Human Hippocampus Supports the Formation of New Episodic Memories. Cerebral Cortex. 2014;26(1):268–278. doi:10.1093/cercor/bhu232.

10. Goutagny R, Gu N, Cavanagh C, Jackson J, Chabot J, Quirion R, et al. Alterations in hippocampal network oscillations and theta–gamma coupling arise before Aβ overproduction in a mouse model of Alzheimer’s disease. European Journal of Neuroscience. 2013;37(12):1896–1902. doi:10.1111/ejn.12233.

11. Bazzigaluppi P, Beckett TL, Koletar MM, Lai AY, Joo IL, Brown ME, et al. Early-stage attenuation of phase-amplitude coupling in the hippocampus and medial prefrontal cortex in a transgenic rat model of Alzheimer’s disease. Journal of Neurochemistry. 2017;144(5):669–679. doi:10.1111/jnc.14136.

12. Kitchigina VF. Alterations of Coherent Theta and Gamma Network Oscillations as an Early Biomarker of Temporal Lobe Epilepsy and Alzheimer’s Disease. Frontiers in Integrative Neuroscience. 2018;12. doi:10.3389/fnint.2018.00036.

13. van den Berg M, Toen D, Verhoye M, Keliris GA. Alterations in theta-gamma coupling and sharp wave-ripple, signs of prodromal hippocampal network impairment in the TgF344-AD rat model. Frontiers in Aging Neuroscience. 2023;15. doi:10.3389/fnagi.2023.1081058.

14. Suthana N, Haneef Z, Stern J, Mukamel R, Behnke E, Knowlton B, et al. Memory Enhancement and Deep-Brain Stimulation of the Entorhinal Area. New England Journal of Medicine. 2012;366(6):502–510. doi:10.1056/nejmoa1107212.

15. Titiz AS, Hill MRH, Mankin EA M Aghajan Z, Eliashiv D, Tchemodanov N, et al. Theta-burst microstimulation in the human entorhinal area improves memory specificity. eLife. 2017;6. doi:10.7554/elife.29515.

16. Jun S, Kim JS, Chung CK. Direct Stimulation of Human Hippocampus During Verbal Associative Encoding Enhances Subsequent Memory Recollection. Frontiers in Human Neuroscience. 2019;13. doi:10.3389/fnhum.2019.00023.

17. Jacobs J, Miller J, Lee SA, Coffey T, Watrous AJ, Sperling MR, et al. Direct Electrical Stimulation of the Human Entorhinal Region and Hippocampus Impairs Memory. Neuron. 2016;92(5):983–990. doi:10.1016/j.neuron.2016.10.062.

18. Merkow MB, Burke JF, Ramayya AG, Sharan AD, Sperling MR, Kahana MJ. Stimulation of the human medial temporal lobe between learning and recall selectively enhances forgetting. Brain Stimulation. 2017;10(3):645–650. doi:10.1016/j.brs.2016.12.011.

19. Hansen N, Chaieb L, Derner M, Hampel KG, Elger CE, Surges R, et al. Memory encoding-related anterior hippocampal potentials are modulated by deep brain stimulation of the entorhinal area. Hippocampus. 2017;28(1):12–17. doi:10.1002/hipo.22808.

20. Hampson RE, Song D, Robinson BS, Fetterhoff D, Dakos AS, Roeder BM, et al. Developing a hippocampal neural prosthetic to facilitate human memory encoding and recall. Journal of Neural Engineering. 2018;15(3):036014. doi:10.1088/1741-2552/aaaed7.

21. Ezzyat Y, Wanda PA, Levy DF, Kadel A, Aka A, Pedisich I, et al. Closed-loop stimulation of temporal cortex rescues functional networks and improves memory. Nature Communications. 2018;9(1). doi:10.1038/s41467-017-02753-0.

22. Khan IS, D’Agostino EN, Calnan DR, Lee JE, Aronson JP. Deep Brain Stimulation for Memory Modulation: A New Frontier. World Neurosurgery. 2019;126:638–646. doi:10.1016/j.wneu.2018.12.184.

23. Mankin EA, Fried I. Modulation of Human Memory by Deep Brain Stimulation of the Entorhinal-Hippocampal Circuitry. Neuron. 2020;106(2):218–235. doi:10.1016/j.neuron.2020.02.024.

24. Gupta A, Vardalakis N, Wagner FB. Neuroprosthetics: from sensorimotor to cognitive disorders. Communications Biology. 2023;6(1). doi:10.1038/s42003-022-04390-w.

25. Mohan UR, Jacobs J. Why does invasive brain stimulation sometimes improve memory and sometimes impair it? PLOS Biology. 2024;22(10):e3002894. doi:10.1371/journal.pbio.3002894.

26. Segneri M, Bi H, Olmi S, Torcini A. Theta-Nested Gamma Oscillations in Next Generation Neural Mass Models. Frontiers in Computational Neuroscience. 2020;14. doi:10.3389/fncom.2020.00047.

27. Sengupta S, Talidou A, Lefebvre J, Skinner FK. Cell-type-specific contributions to theta-gamma coupled rhythms in the hippocampus. Network Neuroscience. 2025;9(1):100–124. doi:10.1162/netn_a_00427.

28. Bezaire MJ, Raikov I, Burk K, Vyas D, Soltesz I. Interneuronal mechanisms of hippocampal theta oscillations in a full-scale model of the rodent CA1 circuit. eLife. 2016;5. doi:10.7554/elife.18566.

29. Ponzi A, Dura-Bernal S, Migliore M. Theta-gamma phase amplitude coupling in a hippocampal CA1 microcircuit. PLOS Computational Biology. 2023;19(3):e1010942. doi:10.1371/journal.pcbi.1010942.

30. Aussel A, Buhry L, Tyvaert L, Ranta R. A detailed anatomical and mathematical model of the hippocampal formation for the generation of sharp-wave ripples and theta-nested gamma oscillations. Journal of Computational Neuroscience. 2018;45(3):207–221. doi:10.1007/s10827-018-0704-x.

31. Mysin I. A Model of the CA1 Field Rhythms. eneuro. 2021;8(6):ENEURO.0192–21.2021. doi:10.1523/eneuro.0192-21.2021.

32. Vardalakis N, Aussel A, Rougier NP, Wagner FB. A dynamical computational model of theta generation in hippocampal circuits to study theta-gamma oscillations during neurostimulation. eLife. 2024;12. doi:10.7554/elife.87356.

33. Rattay F. The basic mechanism for the electrical stimulation of the nervous system. Neuroscience. 1999;89(2):335–346. doi:10.1016/s0306-4522(98)00330-3.

34. Rattay F, Resatz S, Lutter P, Minassian K, Jilge B, Dimitrijevic MR. Mechanisms of Electrical Stimulation with Neural Prostheses. Neuromodulation: Technology at the Neural Interface. 2003;6(1):42–56. doi:10.1046/j.1525-1403.2003.03006.x.

35. Bingham CS, Loizos K, Yu GJ, Gilbert A, Bouteiller JMC, Song D, et al. Model-Based Analysis of Electrode Placement and Pulse Amplitude for Hippocampal Stimulation. IEEE Transactions on Biomedical Engineering. 2018;65(10):2278–2289. doi:10.1109/tbme.2018.2791860.

36. Farzad S, Wei T, Bouteiller JMC, Lazzi G. Dentate gyrus granule cell activation following extracellular electrical stimulation: a multi-scale computational model to guide hippocampal neurostimulation strategies. Frontiers in Computational Neuroscience. 2025;19. doi:10.3389/fncom.2025.1638002.

37. Grill WM. Modeling the effects of electric fields on nerve fibers: influence of tissue electrical properties. IEEE Transactions on Biomedical Engineering. 1999;46(8):918–928. doi:10.1109/10.775401.

38. McIntyre CC, Richardson AG, Grill WM. Modeling the Excitability of Mammalian Nerve Fibers: Influence of Afterpotentials on the Recovery Cycle. Journal of Neurophysiology. 2002;87(2):995–1006. doi:10.1152/jn.00353.2001.

39. Zhang TC, Grill WM. Modeling deep brain stimulation: point source approximation versus realistic representation of the electrode. Journal of Neural Engineering. 2010;7(6):066009. doi:10.1088/1741-2560/7/6/066009.

40. Klausberger T, Somogyi P. Neuronal Diversity and Temporal Dynamics: The Unity of Hippocampal Circuit Operations. Science. 2008;321(5885):53–57. doi:10.1126/science.1149381.

41. Booker SA, Vida I. Morphological diversity and connectivity of hippocampal interneurons. Cell and Tissue Research. 2018;373(3):619–641. doi:10.1007/s00441-018-2882-2.

42. Neymotin SA, Lazarewicz MT, Sherif M, Contreras D, Finkel LH, Lytton WW. Ketamine Disrupts Theta Modulation of Gamma in a Computer Model of Hippocampus. The Journal of Neuroscience. 2011;31(32):11733–11743. doi:10.1523/jneurosci.0501-11.2011.

43. Antonoudiou P, Tan YL, Kontou G, Upton AL, Mann EO. Parvalbumin and Somatostatin Interneurons Contribute to the Generation of Hippocampal Gamma Oscillations. The Journal of Neuroscience. 2020;40(40):7668–7687. doi:10.1523/jneurosci.0261-20.2020.

44. McIntyre CC, Grill WM. Extracellular Stimulation of Central Neurons: Influence of Stimulus Waveform and Frequency on Neuronal Output. Journal of Neurophysiology. 2002;88(4):1592–1604. doi:10.1152/jn.2002.88.4.1592.

45. McIntyre CC, Grill WM, Sherman DL, Thakor NV. Cellular Effects of Deep Brain Stimulation: Model-Based Analysis of Activation and Inhibition. Journal of Neurophysiology. 2004;91(4):1457–1469. doi:10.1152/jn.00989.2003.

46. Meier S, Bräuer AU, Heimrich B, Nitsch R, Savaskan NE. Myelination in the hippocampus during development and following lesion. Cellular and Molecular Life Sciences (CMLS). 2004;61(9):1082–1094. doi:10.1007/s00018-004-3469-5.

47. Nickel M, Gu C. Regulation of Central Nervous System Myelination in Higher Brain Functions. Neural Plasticity. 2018;2018:1–12. doi:10.1155/2018/6436453.

48. Csicsvari J, Hirase H, Czurkó A, Mamiya A, Buzsáki G. Oscillatory Coupling of Hippocampal Pyramidal Cells and Interneurons in the Behaving Rat. The Journal of Neuroscience. 1999;19(1):274–287. doi:10.1523/jneurosci.19-01-00274.1999.

49. Brocker DT, Grill WM. 1. In: Principles of electrical stimulation of neural tissue. Elsevier; 2013. p. 3–18. Available from: http://dx.doi.org/10.1016/B978-0-444-53497-2.00001-2.

50. McIntyre CC, Grill WM. Extracellular Stimulation of Central Neurons: Influence of Stimulus Waveform and Frequency on Neuronal Output. Journal of Neurophysiology. 2002;88(4):1592–1604. doi:10.1152/jn.2002.88.4.1592.

51. McIntyre CC, Savasta M, Kerkerian-Le Goff L, Vitek JL. Uncovering the mechanism(s) of action of deep brain stimulation: activation, inhibition, or both. Clinical Neurophysiology. 2004;115(6):1239–1248. doi:10.1016/j.clinph.2003.12.024.

52. Romaro C, Najman FA, Lytton WW, Roque AC, Dura-Bernal S. NetPyNE Implementation and Scaling of the Potjans-Diesmann Cortical Microcircuit Model. Neural Computation. 2021;33(7):1993–2032. doi:10.1162/neco_a_01400.

53. McIntyre CC, Grill WM. Excitation of Central Nervous System Neurons by Nonuniform Electric Fields. Biophysical Journal. 1999;76(2):878–888. doi:10.1016/s0006-3495(99)77251-6.

54. Kringelbach ML, Jenkinson N, Owen SLF, Aziz TZ. Translational principles of deep brain stimulation. Nature Reviews Neuroscience. 2007;8(8):623–635. doi:10.1038/nrn2196.

55. McIntyre CC, Anderson RW. Deep brain stimulation mechanisms: the control of network activity via neurochemistry modulation. Journal of Neurochemistry. 2016;139(S1):338–345. doi:10.1111/jnc.13649.

56. Benazzouz A, Hamani C. In: Mechanisms of Deep Brain Stimulation. Springer International Publishing; 2020. p. 29–37. Available from: http://dx.doi.org/10.1007/978-3-030-36346-8_3.

57. Buzsáki G. Theta Oscillations in the Hippocampus. Neuron. 2002;33(3):325–340. doi:10.1016/s0896-6273(02)00586-x.

58. Colgin LL. Mechanisms and Functions of Theta Rhythms. Annual Review of Neuroscience. 2013;36(1):295–312. doi:10.1146/annurev-neuro-062012-170330.

59. Mysin IE, Kitchigina VF, Kazanovich YB. Phase relations of theta oscillations in a computer model of the hippocampal CA1 field: Key role of Schaffer collaterals. Neural Networks. 2019;116:119–138. doi:10.1016/j.neunet.2019.04.004.

60. Anderson TR, Hu B, Iremonger K, Kiss ZHT. Selective Attenuation of Afferent Synaptic Transmission as a Mechanism of Thalamic Deep Brain Stimulation-Induced Tremor Arrest. The Journal of Neuroscience. 2006;26(3):841–850. doi:10.1523/jneurosci.3523-05.2006.

61. Rosenbaum R, Zimnik A, Zheng F, Turner RS, Alzheimer C, Doiron B, et al. Axonal and synaptic failure suppress the transfer of firing rate oscillations, synchrony and information during high frequency deep brain stimulation. Neurobiology of Disease. 2014;62:86–99. doi:10.1016/j.nbd.2013.09.006.

62. Hodgkin AL, Huxley AF. A quantitative description of membrane current and its application to conduction and excitation in nerve. The Journal of Physiology. 1952;117(4):500–544. doi:10.1113/jphysiol.1952.sp004764.

63. Hines M. NEURON and Python. Frontiers in Neuroinformatics. 2009;3. doi:10.3389/neuro.11.001.2009.

64. West MJ, Coleman PD, Flood DG, Troncoso JC. Differences in the pattern of hippocampal neuronal loss in normal ageing and Alzheimer’s disease. The Lancet. 1994;344(8925):769–772. doi:10.1016/s0140-6736(94)92338-8.

65. Bezaire MJ, Soltesz I. Quantitative assessment of CA1 local circuits: Knowledge base for interneuron-pyramidal cell connectivity: Quantitative Assessment Of Ca1 Local Circuits. Hippocampus. 2013;23(9):751–785. doi:10.1002/hipo.22141.

66. The Hippocampus Book. Oxford University Press; 2006. Available from: http://dx.doi.org/10.1093/acprof:oso/9780195100273.001.0001.

67. Cutsuridis V, Cobb S, Graham BP. Encoding and retrieval in a model of the hippocampal CA1 microcircuit. Hippocampus. 2009;20(3):423–446. doi:10.1002/hipo.20661.

68. Wheeler DW, Kopsick JD, Sutton N, Tecuatl C, Komendantov AO, Nadella K, et al. Hippocampome.org 2.0 is a knowledge base enabling data-driven spiking neural network simulations of rodent hippocampal circuits. eLife. 2024;12. doi:10.7554/elife.90597.

69. Hines ML, Carnevale NT. The NEURON Simulation Environment. Neural Computation. 1997;9(6):1179–1209. doi:10.1162/neco.1997.9.6.1179.

